# IgG4^+^ plasma cell enrichment and λ-chain-biased BCR remodeling drive low-grade autoimmunity in chronic obstructive pulmonary disease

**DOI:** 10.64898/2026.06.25.734436

**Authors:** Luo Duan, Han Zhao, Yanqi Xia, Xiaoxia Ren, Huanyu Long, Lun Li, Mi Mu, Zengqing Liu, Kaixin Li, Jie Liu, Yunpeng Dou, Ye Cui, Yan Chen, Zhe Lv, Chris J Corrigan, Sebastian Lennox Johnston, Wei Wang, Huihui Yuan, Ying Sun

**Affiliations:** Department of Immunology, School of Basic Medical Sciences, Capital Medical University, Beijing, China; Beijing Shijitan Hospital, Capital Medical University, Beijing, China; Immune Dysfunction and Pulmonary Fibrosis Joint Laboratory for Clinical Medicine, Capital Medical University, Beijing, China; Department of Respiratory and Critical Care Medicine, The Eighth Medical Center of Chinese PLA General Hospital, Beijing, China; Department of Respiratory Medicine, Capital Medical University, Beijing, China; Department of Pulmonary and Critical Care Medicine, Peking University Third Hospital, Beijing, China; Department of Allergy, Beijing Key Laboratory of Precision Medicine for Diagnosis and Treatment on Allergic Diseases, Peking Union Medical College Hospital, Chinese Academy of Medical Sciences & Peking Union Medical College, Beijing, China; Non-Invasive Diagnosis and Immunotherapy Laboratory for Rheumatic Immune Diseases, Capital Medical University, Beijing, China; Faculty of Life Sciences & Medicine, School of Immunology & Microbial Sciences, King’s College London, London, UK; National Heart and Lung Institute, Faculty of Medicine, Imperial College London, London, UK

**Keywords:** Chronic obstructive pulmonary disease (COPD), B cell, Autoimmunity, IgG4, Transitional B cell

## Abstract

**Background:** This study aimed to elucidate B cell subset pathology in COPD, a poorly characterized area, with a focus on its similarities to and differences from classical autoimmune disorders.

**Methods:** The single-cell RNA-sequencing (scRNA-seq) data from COPD and autoimmune diseases were obtained from Gene Expression Omnibus (GEO) for comparative analyses of B cell subsets and functions via differentially expressed genes (DEGs), KEGG, protein-protein interaction (PPI), and cell–cell communication analyses. Serum IgG4 was measured by ELISA and correlated with clinical parameters. The peripheral blood B cells were sorted by flow cytometry for single-cell B cell receptor (BCR) sequencing. A v-Abl/Bcl2 pro-B cell line was stimulated with cigarette smoke extract (CSE) to assess abnormal development *in vitro*.

**Results:** In lung tissue, IgG4^+^ plasma cells were enriched and expressed BCR activation/inflammatory genes and TNF/NF-κB/MAPK pathways. Serum IgG4 concentrations correlated negatively with pre-and post-bronchodilator FEV_1_/FVC. B cell interacted with monocytes, macrophages, fibroblasts and endothelial cells via IL-1β/IL-6, integrin and chemokine signalling, contributing to chronic inflammation and remodelling. In peripheral blood, transitional T1 B cells were increased, accompanied by λ-chain enrichment and increased *IGLV1-47* usage, as well as enrichment of autoimmune pathways. In the bone marrow, the numbers of pre-B I cells were increased while those of small pre-B III cells were reduced, with altered expression of BCR development genes. CSE stimulation of the pro-B cell line reduced λ5 expression in a concentration-dependent manner.

**Conclusions:** The autoimmune abnormalities in COPD appear more restricted, although IgG4 antibody generation may contribute to immune-mediated lung damage.

## 1. Introduction

Chronic obstructive pulmonary disease (COPD) is the third leading cause of death worldwide. It is a chronic inflammatory airways disease characterised by persistent airflow limitation, primarily driven by prolonged exposure to noxious inhaled particles, most notably tobacco smoke and air pollutants, alongside contributions from genetic susceptibility, developmental factors and social determinants of health [1,2]. Despite its high prevalence, the precise mechanisms underlying COPD pathogenesis remain incompletely elucidated.

In 2003, Bartolomé Celli and colleagues first proposed the autoimmune hypothesis of COPD. This hypothesis was based on observations of increased pulmonary immune cellular infiltration, persistent airways inflammation despite smoking cessation, a higher prevalence of autoimmune diseases such as rheumatoid arthritis (RA) among smokers, and the presence of antinuclear antibodies (ANAs) in patients with COPD [3]. Since then, accumulating evidence has substantiated the involvement of autoimmunity in COPD pathogenesis. For instance, protein microarray analyses profiling over 19,000 human proteins have revealed distinct autoantibody signatures in smokers, both with and without COPD [4]. Furthermore, integrative omics analyses have implicated B-cell receptor (BCR) signalling in the emphysema phenotype of COPD [5]. Notably, emphysema severity correlates with genes involved in B-cell maturation and antibody production, while patients with severe emphysema exhibit chronic B-cell activation and enrichment of autoimmune-related pathways [6]. Collectively, these findings suggest that B-cell autoimmune abnormalities may contribute to disease progression, particularly in a subset of patients with emphysema-dominant COPD.

COPD shares several immunological hallmarks with classical autoimmune diseases, including autoantigen exposure, interferon activation and B-cell dysregulation. Certain clinical and immunological features of COPD closely mirror those observed in systemic autoimmune disorders. For example, protein citrullination, a hallmark autoantigenic process in RA, is upregulated in COPD lung tissue irrespective of smoking status [7]. Similarly, cigarette smoke exposure in mice induces the release of extracellular double-stranded DNA (dsDNA), triggering inflammatory responses and ANAs production reminiscent of systemic lupus erythematosus (SLE) [8]. A large-scale clinical study has further demonstrated overlapping blood transcriptomic signatures between stable COPD and SLE, suggesting a persistent, interferon-related inflammatory program associated with immune-mediated lung injury [9]. In addition, patients with COPD display an imbalanced ratio of effector B-cells (Beff) to regulatory B-cells (Breg), alongside increased memory B cells and enhanced plasma-cell-related gene expression, abnormalities that parallel those observed in systemic sclerosis (SSc) [10–12].

Given the emerging pathogenic role of B cells in COPD, B-cell-targeted therapies have garnered increasing attention. Although rituximab (anti-CD20) was evaluated in clinical trials for therapy of diseases such as multiple sclerosis, its development was halted because of an increased risk of pulmonary infection [13]. Preclinical studies in cigarette smoke-exposed mice implicate B cell activating factor (BAFF) in the production of cigarette smoke-induced pulmonary autoantibody production, suggesting that BAFF blockade may have therapeutic potential in COPD [14]. Moreover, data suggest that inebilizumab (MEDI-551, anti-CD19) which targets plasma cells may be effective for the treatment of emphysema along with a range of autoimmune diseases [11]. Together, these findings underscore the therapeutic relevance of B-cell-targeted strategies in selected COPD populations exhibiting aberrant B-cell responses.

Despite the growing body of evidence implicating B cells in COPD, the specific nature of B cell autoimmune abnormalities in this disease remain substantially less well characterised than in classical autoimmune diseases. It is clear that further investigation is warranted to identify the key B-cell subsets, core regulatory genes and associated pathways driving COPD pathogenesis, as well as to delineate the similarities and distinctions between COPD and other autoimmune diseases.

With the rapid advancement of sequencing technologies and bioinformatics, large-scale characterisation of immune-cell heterogeneity has become increasingly feasible. In the present study, we integrated single-cell transcriptomic analyses of lung tissue, peripheral blood and bone marrow from patients with, or murine models with COPD and autoimmune diseases, utilizing publicly available datasets from the Gene

Expression Omnibus (GEO) database. These bioinformatic findings were further validated through serum IgG4 quantification, single-cell BCR sequencing of sorted peripheral blood B cells, and *in vitro* cigarette smoke extract (CSE) stimulation experiments using a v-Abl/Bcl2 pro-B cell line. Our study provides multi-dimensional evidence supporting the involvement of B-cell autoimmunity and aberrant development in COPD pathogenesis.

## 2. Materials and Methods

### 2.1 Ethics approval

All human studies were approved by the Institutional Review Board of Capital Medical University (approval no. Z2021SY025). Written informed consent was obtained from all participants or their legal guardians prior to enrolment. COPD was diagnosed in accordance with the 2023 Global Initiative for Chronic Obstructive Lung Disease (GOLD) criteria, defined by a post-bronchodilator forced expiratory volume in one second/forced vital capacity (FEV_1_/FVC) ratio <0.70. Diagnosis also required the presence of compatible respiratory symptoms (e.g., dyspnoea and chronic cough) and relevant exposure history, including a smoking history of ≥10 pack-years. Patients with a prior diagnosis of asthma or other chronic respiratory diseases were excluded from the study [15].

### 2.2 Demographic and clinical data collection

For IgG4 detection and correlation analysis, clinical data collected for each patient with COPD included age, sex, body mass index (BMI), and smoking-related variables such as cigarettes smoked per day (CPD), smoking duration, smoking cessation status, and years since cessation. Pulmonary function parameters included post-bronchodilator FEV_1_, FVC and FEV_1_/FVC ratio. Symptom burden and disease severity were assessed using the modified Medical Research Council (mMRC) dyspnoea scale, the COPD Assessment Test (CAT), GOLD grade (based on predicted FEV_1_%) and the GOLD ABCD classification (2023–2026 criteria). We also recorded the number of acute exacerbations within the previous 12 months. Routine haematological parameters included white blood cell count, neutrophil, lymphocyte, monocyte, eosinophil and basophil counts, neutrophil-to-lymphocyte ratio (NLR), platelet count, haemoglobin level and haematocrit (Table 1).

**Table 1.**
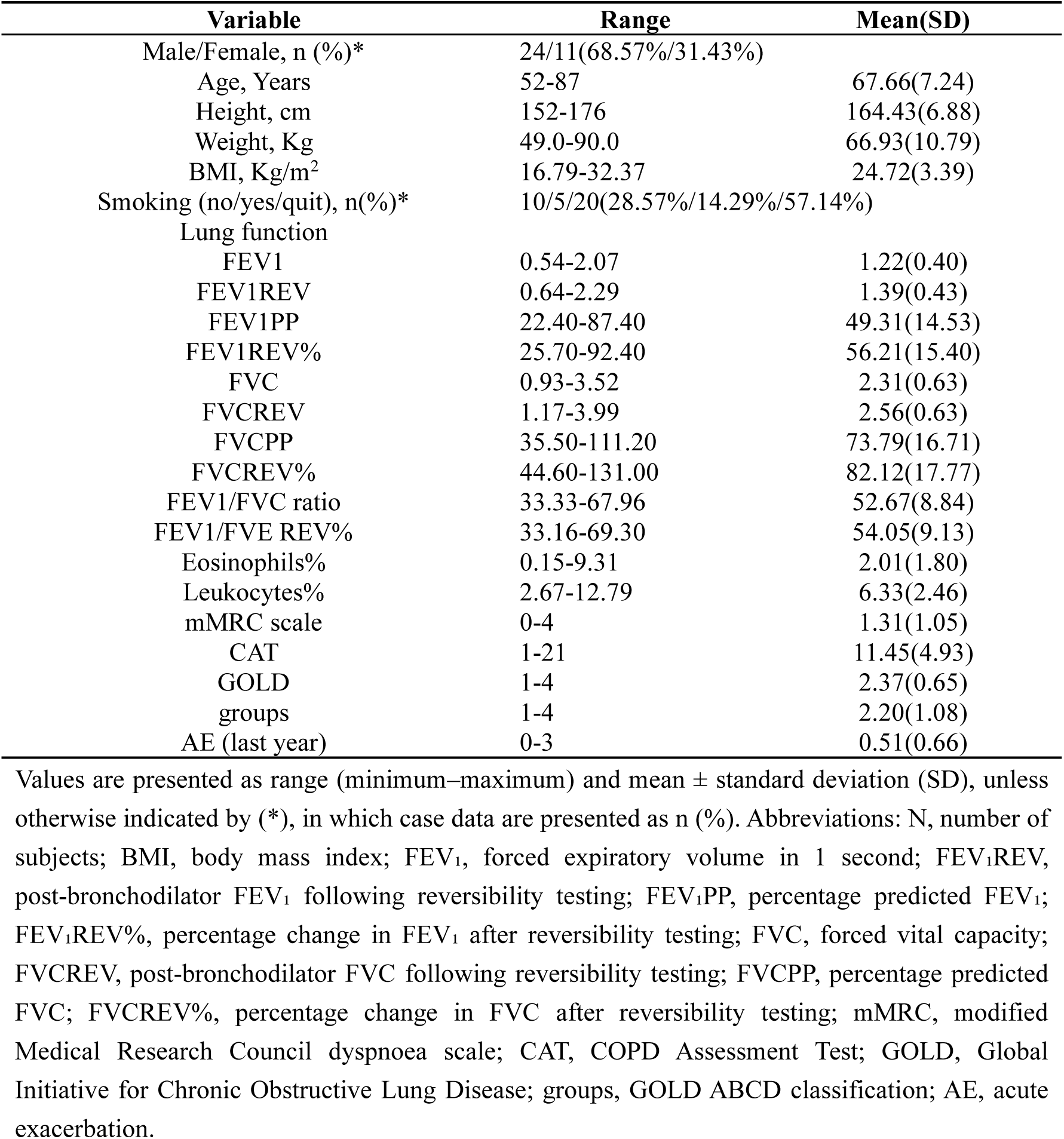
Demographic, clinical, and functional characteristics of COPD patients (n = 35)

For single-cell BCR sequencing, demographic and clinical information for COPD patients and controls included sex, age, height, weight, BMI, tobacco load, smoking duration, FEV_1_/FVC ratio, mMRC scale, and CAT (Table 2).

**Table 2.** Baseline characteristics of COPD patients and healthy controls.

| **Variable** | **HC group (n=4)** | **COPD group (n=3)** | **P** |
| --- | --- | --- | --- |
| Male/Female, n (%)* | 3/1(75%/25%) | 3/0(100%/0%) | 0.052 |
| Age, Years | 57.75±14.22 | 56.00±12.77 | 0.545 |
| Height, m | 1.70±0.07 | 1.74±0.02 | 0.155 |
| Weight, Kg | 73.63±13.39 | 83.67±18.04 | 0.648 |
| BMI, Kg/m<sup>2</sup> | 25.44±3.38 | 27.63±5.96 | 0.345 |
| Tobacco load (pack/year) | 197.75±152.03 | 324.33±70.44 | 0.382 |
| Smoking (years) | 27.50±22.17 | 38.67±22.59 | 0.049 |
| FEV1/FVC ratio | 76.50±3.42 | 58.31±16.83 | 0.083 |
| mMRC scale | 0±0.00 | 0.33±0.58 | 0.005 |
| CAT | 0±0.00 | 15.33±8.50 | 0.008 |

Values are presented as mean ± standard deviation (SD), unless otherwise indicated by (\*), in which case data are presented as n (%). Abbreviations: BMI, body mass index; FEV<sub>1</sub>, forced expiratory volume in 1 second; FVC, forced vital capacity; mMRC, modified Medical Research Council dyspnoea scale; CAT, COPD Assessment Test.

### 2.3 Human PBMC collection and cell isolation

Peripheral blood mononuclear cells (PBMCs) were isolated using HISTOPAQUE-1077 (Sigma-Aldrich) according to the manufacturer’s instructions. Briefly, 2 mL of fresh peripheral blood collected in EDTA anticoagulant tubes were layered onto HISTOPAQUE-1077 and centrifuged. The mononuclear cell layer at the plasma–HISTOPAQUE interface was collected, transferred to a new tube and washed twice with phosphate-buffered saline (PBS; Gibco). Cells were subsequently treated with red blood cell lysis buffer (Miltenyi Biotec) on ice for 1–2 min, washed twice, and resuspended in fluorescence-activated cell sorting (FACS) buffer consisting of PBS supplemented with 2% foetal bovine serum (FBS) at a final concentration of approximately 1 × 10⁶ cells/mL.

### 2.4 FACS for B-cell sorting

For live B-cell sorting, PBMCs were stained with anti-CD45-FITC (BioLegend, clone HI30), anti-CD19-PerCP-Cy5.5 (BioLegend, clone HIB19) and fixable viability stain 506 (FVS506; BD Horizon™) for 30 min on ice in the dark. Following staining, cells were washed twice and resuspended in MACSQuant Tyto running buffer (Miltenyi Biotec) at a concentration of 2–4 × 10⁶ cells/mL. Cell sorting was performed using a MACSQuant Tyto cell sorter (Miltenyi Biotec) equipped with 405 nm, 488 nm, and 561 nm lasers and a closed cartridge-based system. Sorted cells with a purity ≥90% were collected into the positive chamber, transferred into 1.5 mL centrifuge tubes using gel-loading tips and maintained on ice prior to single-cell BCR sequencing. All procedures were performed under sterile conditions at 4°C.

### 2.5 Single-cell 5′ BCR sequencing

Immediately after sorting, live CD45^+^CD19^+^ B cells were centrifuged (300 × g, 5 min, 4°C) and resuspended in PBS containing 0.04% bovine serum albumin (BSA) at a concentration of 700-1,200 cells/μL. Single-cell libraries were generated using the Chromium Next GEM Single Cell 5′ Reagent Kit v2 (10x Genomics, PN-1000265) according to the manufacturer’s protocol. For BCR libraries, nested PCR amplification of immunoglobulin heavy-chain (IgH) and light-chain (Igκ and Igλ) transcripts was performed from 5′ barcoded cDNA following the 10x Genomics V(D)J workflow. Final libraries were quantified using a Qubit 4.0 fluorometer (Thermo Fisher Scientific), and fragment size distributions were evaluated using a Bioanalyzer 2100 (Agilent Technologies) with a High Sensitivity DNA chip. Paired-end sequencing was conducted on an Illumina NovaSeq X Plus platform using the Xp workflow, targeting a minimum sequencing depth of 5,000 read pairs per cell for BCR libraries. FASTQ files were processed using Cell Ranger v7.1 (10x Genomics) with the Cell Ranger vdj Pipeline aligned to the GRCh38 reference genome (refdata-gex-GRCh38-2020-A). Productive and non-productive V(D)J rearrangements were identified for each cell.

### 2.6 Single-cell RNA-seq data collection from GEO

Single-cell RNA sequencing (scRNA-seq) datasets for COPD and autoimmune diseases were obtained from the GEO database. Detailed accession numbers and sample information are provided in Supplementary Table 1.

### 2.7 Quality control of scRNA-seq data

Quality control was performed using Seurat (v4.4.0) to remove doublets, damaged cells and dead or dying cells prior to data integration and batch correction [16]. The Harmony package (v0.1.0) was used for sample integration and reduction of batch effects. For publicly available scRNA-seq datasets, cells expressing fewer than 150 genes or more than 5,000 genes, as well as cells with >10% mitochondrial gene expression were excluded. For single-cell 5′ BCR sequencing datasets, cells expressing fewer than 200 genes or more than 2,500 genes, together with those with >10% mitochondrial gene expression were excluded. Data normalization and integration were performed using the NormalizeData function following the standard Seurat workflow. Gene expression values were log-transformed and subsequently scaled at the gene level using the ScaleData function.

### 2.8 Cluster annotation and quantification

Principal component analysis (PCA) was performed using the top 2,000 variable features. Depending on the dataset, the top 5–17 principal components were selected for downstream analyses. Cell clustering was performed in Seurat using a resolution range of 0.2–0.5. Clusters were visualised using Uniform Manifold Approximation and Projection (UMAP) and annotated based on the top five highly expressed genes identified by the FindAllMarkers function, followed by manual verification using CellMarker 2.0 and previously published marker gene references [17–20]. Cell numbers and proportions were subsequently calculated for each cluster.

### 2.9 Differential gene expression analysis

Differentially expressed genes (DEGs) for each cluster were identified relative to all remaining cells using the FindMarkers function in Seurat. The minimum percentage threshold for gene expression within each cluster was set at 0.1. Differential expression between disease and control groups was assessed using the Wilcoxon rank-sum test with thresholds of log₂ fold change >0.25 and p <0.05. For IgG4^+^ plasma cells specifically identified in the COPD group, the FindAllMarkers function was used to identify genes enriched relative to other cell populations applying thresholds of log₂ fold change >0.25 and adjusted p <0.05.

### 2.10 Pathway analysis

Kyoto Encyclopedia of Genes and Genomes (KEGG) pathway enrichment analysis of upregulated and downregulated genes was performed using the R packages clusterProfiler and enrichplot [21]. Significance thresholds were set at p <0.05 for transitional B-cell analyses and adjusted p <0.05 for all other analyses.

### 2.11 Protein–protein interaction (PPI) analysis

DEGs were mapped to the STRING database to construct PPI networks using a confidence score threshold ≥0.400. For hub gene visualisation, interaction data containing log₂ fold change information were imported into Cytoscape (v3.10.0) and analysed using the cytoHubba plugin [22].

### 2.12 Cell–cell communication analysis

Cell–cell communication between B-cell subclusters and other cell populations was analysed using the iTALK package, which evaluates cytokine, growth factor, immune checkpoint and additional receptor–ligand interactions [23]. Receptor–ligand pairs were identified using the STRING database and selected through the FindLR function.

### 2.13 BCR analysis

Cells containing at least one productive heavy-chain rearrangement and one productive light-chain rearrangement (κ or λ) were retained for analysis. Using the R package scRepertoire (v1.6), we evaluated clonal diversity (Shannon entropy and inverse Simpson index), clonal expansion (defined as the proportion of cells belonging to expanded clones containing >1 cell), V(D)J gene usage, somatic hypermutation frequency and immunoglobulin isotype distribution, including IgM, IgD, IgG1–4, IgA1–2, and IgE.

### 2.14 ELISA detection of serum IgG4

Peripheral blood samples were collected, and serum was isolated and stored at −80°C until analysis. Prior to ELISA, serum samples were thawed on ice and centrifuged at 10,000 × g for 5 min at 4°C to remove cryoprecipitates and cellular debris. Serum IgG4 concentrations were measured using a commercially available human IgG4 ELISA kit (Thermo Fisher Scientific; cat. no. BMS2095TWO) according to the manufacturer’s instructions.

### 2.15 Correlation analysis

Associations between the proportion of IgG4^+^ plasma cells and clinical characteristics were assessed using Spearman’s rank correlation coefficient. All analyses were performed using GraphPad Prism software (version 10.1). The correlation between the proportion of a given cell type (as the independent variable) and the corresponding clinical parameter (as the dependent variable) was analysed. A two-tailed P < 0.05 was considered statistically significant.

### 2.16 Culture and stimulation

v-Abl/Bcl2 pro-B cell lines were maintained in complete RPMI 1640 medium (Thermo Fisher Scientific) supplemented with 10% FBS, 50 U/mL penicillin, 50 μg/mL streptomycin, 2 mM L-glutamine, 1× MEM non-essential amino acids, 1 mM sodium pyruvate, 50 μM 2-mercaptoethanol (Sigma-Aldrich), and 20 mM HEPES (pH 7.4). Cells were cultured at 37°C in a humidified atmosphere containing 5% CO₂ and maintained at a density between 1 × 10⁶ cells/mL by subculturing every 2–3 days [24].

To synchronise cells in the G1 phase and induce V(D)J recombination, exponentially growing v-Abl/Bcl2 pro-B cells were treated with the Abl kinase inhibitor imatinib (STI-571; 3 μM) for 96 h as a positive control. Experimental groups were treated with CSE at concentrations of 1.25%, 2.5%, 5%, and 10% for 96 h. Following treatment, the expression of BCR-related genes, including λ5, λ5-promoter, Igκ, VpreB-promoter and IL-7R were quantified. Primer sequences are listed in Supplementary Table 2.

Data were analysed and visualized using GraphPad Prism (version 10.1). The relative expression of target genes (normalized to β-actin) was calculated using the 2^−ΔΔCt^ method. Results are presented as the mean ± standard error of the mean (SEM) from four independent experiments (n = 4 per group). One-way analysis of variance (ANOVA) followed by Dunnett’s multiple comparisons test was performed to compare each treatment group with the untreated control group. A P value < 0.05 was considered statistically significant.

### 2.17 Statistical analysis

Unless otherwise specified, all statistical analyses and data visualisation were performed using R (v3.5.2 and v4.0.2). For two-group comparisons, parametric data were analysed using a two-tailed Student’s t-test or Welch’s t-test as appropriate, whereas non-parametric data were analysed using the Wilcoxon signed-rank test. For comparisons involving more than two groups, one-way ANOVA followed by Tukey’s post hoc correction was applied. Data normality was assessed using the shapiro.test function, and homogeneity of variance was evaluated using the leveneTest function from the car package (v3.0-3). Data are presented as the mean ± standard deviation (SD) or the mean ± SEM. A two-sided p value <0.05 was considered statistically significant.

## 3 Results

### 3.1 IgG4^+^ plasma cells are enriched in the lungs of patients with COPD

To characterize B-cell abnormalities in COPD lung tissue, we isolated B cells from integrated single-cell datasets (GSE196638) and performed dimensionality reduction analysis, identifying four major B-cell subclusters (Figure 1A). The overall proportion of B cells was elevated in COPD samples compared with controls (6.72% vs. 3.52%) (Supplementary Figure 1A, C, E). Subclusters were annotated based on top DEGs (Figure 1B) and established markers [25, 26] (Figure 1C). Notably, while memory B cells were increased, IgG1^+^ plasma cells were reduced in COPD lungs (Figure 1D). Strikingly, a distinct IgG4^+^ plasma-cell population, accounting for approximately 5% of total B cells, was identified exclusively in COPD samples (Figure 1A, D).

**Figure 1.**
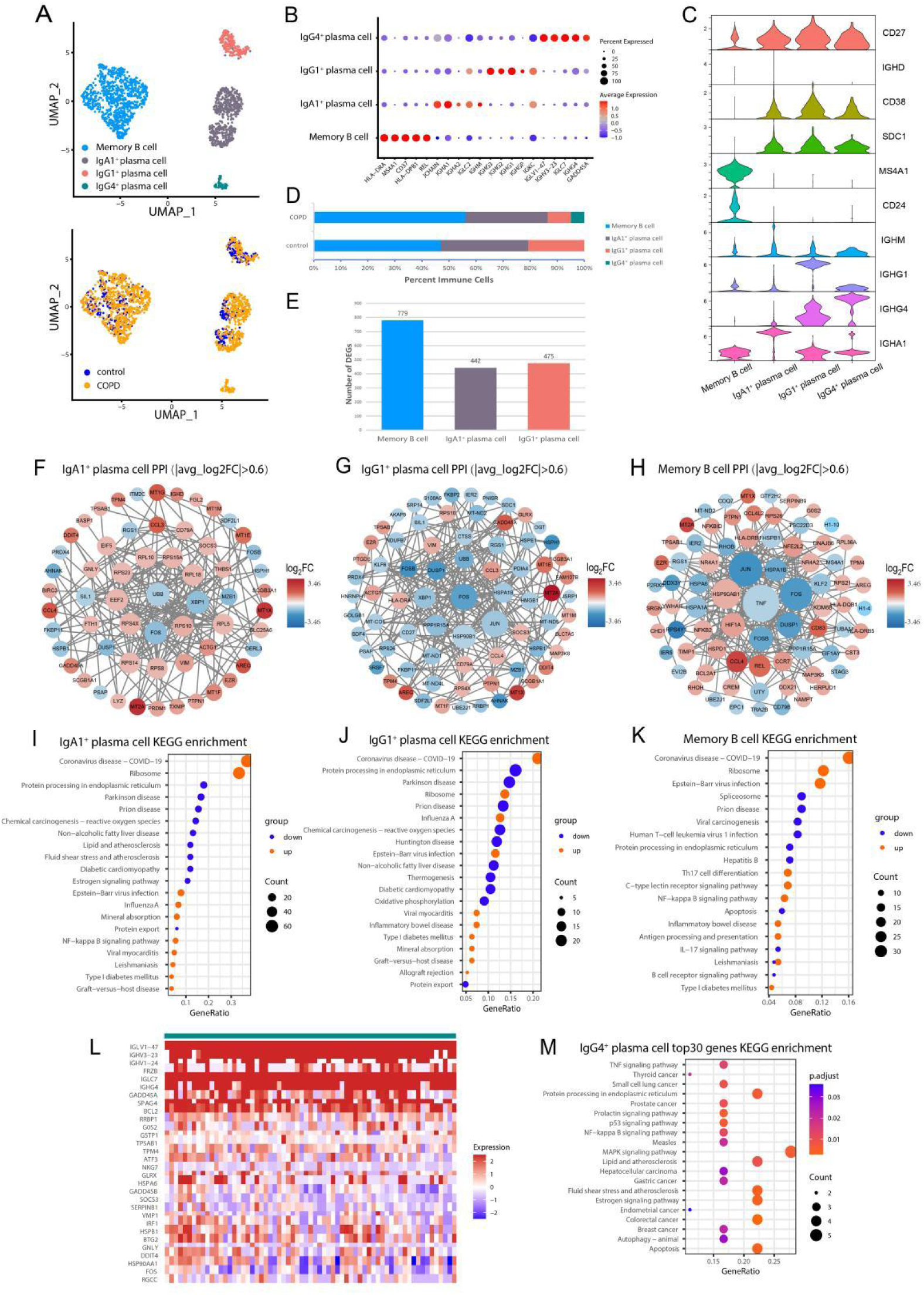
B-cell subclusters in COPD lung tissue. **(A)** Uniform Manifold Approximation and Projection (UMAP) visualization of 1,807 B cells derived from COPD and control lung tissues, coloured according to B-cell subsets and disease groups. **(B)** Dot plot showing the top five marker genes for each B-cell subset. **(C)** Violin plots displaying the expression of representative functional marker genes across B-cell subsets. **(D)** Bar plot showing the relative frequencies of B-cell subsets in COPD and control samples. **(E)** Numbers and proportions of differentially expressed genes (DEGs) identified in each B-cell subset. **(F–H)** Protein–protein interaction (PPI) networks of DEGs (|log₂FC| >0.6) in IgA1^+^ plasma cells, IgG1^+^ plasma cells and memory B cells. Red nodes indicate upregulated genes, whereas blue nodes indicate downregulated genes; node size reflects the number of interacting proteins. **(I–K)** Kyoto Encyclopedia of Genes and Genomes (KEGG) pathway enrichment analyses of upregulated (red) and downregulated (blue) pathways in the corresponding B-cell subsets. **(L)** Heatmap showing the top 30 highly expressed genes in IgG4^+^ plasma cells. **(M)** KEGG pathway enrichment analysis of the top 30 genes identified in IgG4^+^ plasma cells; circle size represents the number of enriched genes within each pathway.

Transcriptional profiling revealed significant alterations in memory B cells, IgA1^+^, and IgG1^+^ plasma cells (Figure 1E). PPI network analysis identified MT2A as consistently upregulated across these B-cell subsets (Figure 1F–H). KEGG enrichment analysis indicated activation of immune and autoimmune-associated pathways, including type 1 diabetes mellitus (T1DM), inflammatory bowel disease (IBD) and Th17 differentiation (Figure 1I–K). Within IgG4^+^ plasma cells, highly expressed genes included BCR activation-related genes (*IGLV1-47, IGHV1-24, IGHV3-23, BCL2,* and *FOS*) and inflammatory-response genes (*HSPA6, HSPB1, HSP90AA1,* and *IRF1*) (Figure 1L). KEGG analysis further exhibited enrichment of TNF, NF-κB, MAPK and p53 signalling pathways (Figure 1M).

Given the enrichment of IgG4^+^ plasma cells in COPD lungs, we quantified serum IgG4 concentrations and correlated them with clinical parameters (Table 1). Serum IgG4 concentrations were associated with age and showed inverse trends with pulmonary function indices, including pre- and post-bronchodilator FEV_1_/FVC ratios (Figure 2A, B, O, T).

**Figure 2.**
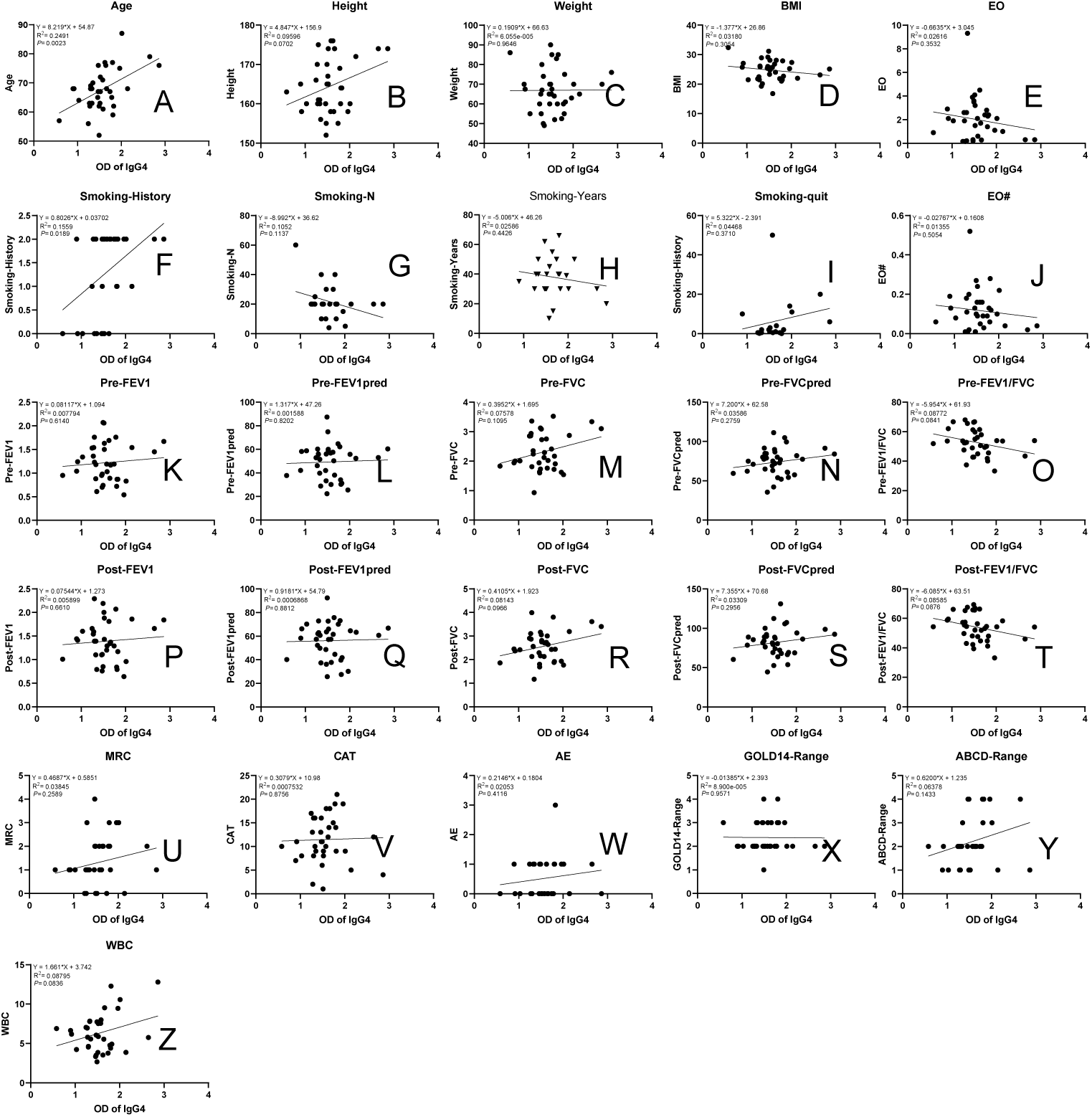
Associations between serum IgG4 concentrations and clinical characteristics in patients with COPD. Spearman correlation analyses between serum IgG4 concentrations and clinical parameters in patients with COPD (n = 35), including age, height, weight, body mass index (BMI), eosinophil count (absolute value and percentage), smoking-related variables (cigarettes per day [CPD], smoking duration, and smoking cessation status), pulmonary function parameters (pre- and post-bronchodilator FEV₁, FEV₁% predicted, FVC, FVC% predicted, and FEV₁/FVC ratio), modified Medical Research Council (mMRC) dyspnoea score, COPD Assessment Test (CAT) score, acute exacerbation (AE) status, GOLD grade, GOLD ABCD classification and white blood cell (WBC) count. Abbreviations: AE, acute exacerbation; BMI, body mass index; CAT, COPD Assessment Test; CPD, cigarettes per day; FEV₁, forced expiratory volume in 1 second; FVC, forced vital capacity; GOLD, Global Initiative for Chronic Obstructive Lung Disease; mMRC, modified Medical Research Council; WBC, white blood cell.

### 3.2 B-cell communication networks are associated with chronic inflammation and tissue remodelling in COPD

Further to investigate the potential interactions between B-cell subsets and the pulmonary microenvironment in COPD, cell–cell communication analyses were performed between B-cell subclusters and immune cells (monocytes, macrophages, NK and T cells) as well as non-immune cells (fibroblasts, endothelial, basal, alveolar epithelial and stromal cells).

Predicted ligand–receptor interactions suggest that IgA1^+^ plasma cells may influence monocyte-macrophage activation through HLA-A/HLA-B–KIR2DL3 signalling and participate in fibroblast and endothelial-cell remodelling through THBS1/TGFB1-associated integrin signalling pathways. Conversely, reduced CCL13-associated signalling implied weaker communication with immune and stromal cell populations. Reciprocal signalling from neighbouring cells was associated with increased expression of ICOSLG, IL1R1, CD44, and TRAF3 on IgA1^+^ plasma cells (Figure 3A–B, Supplementary Table 3).

**Figure 3.**
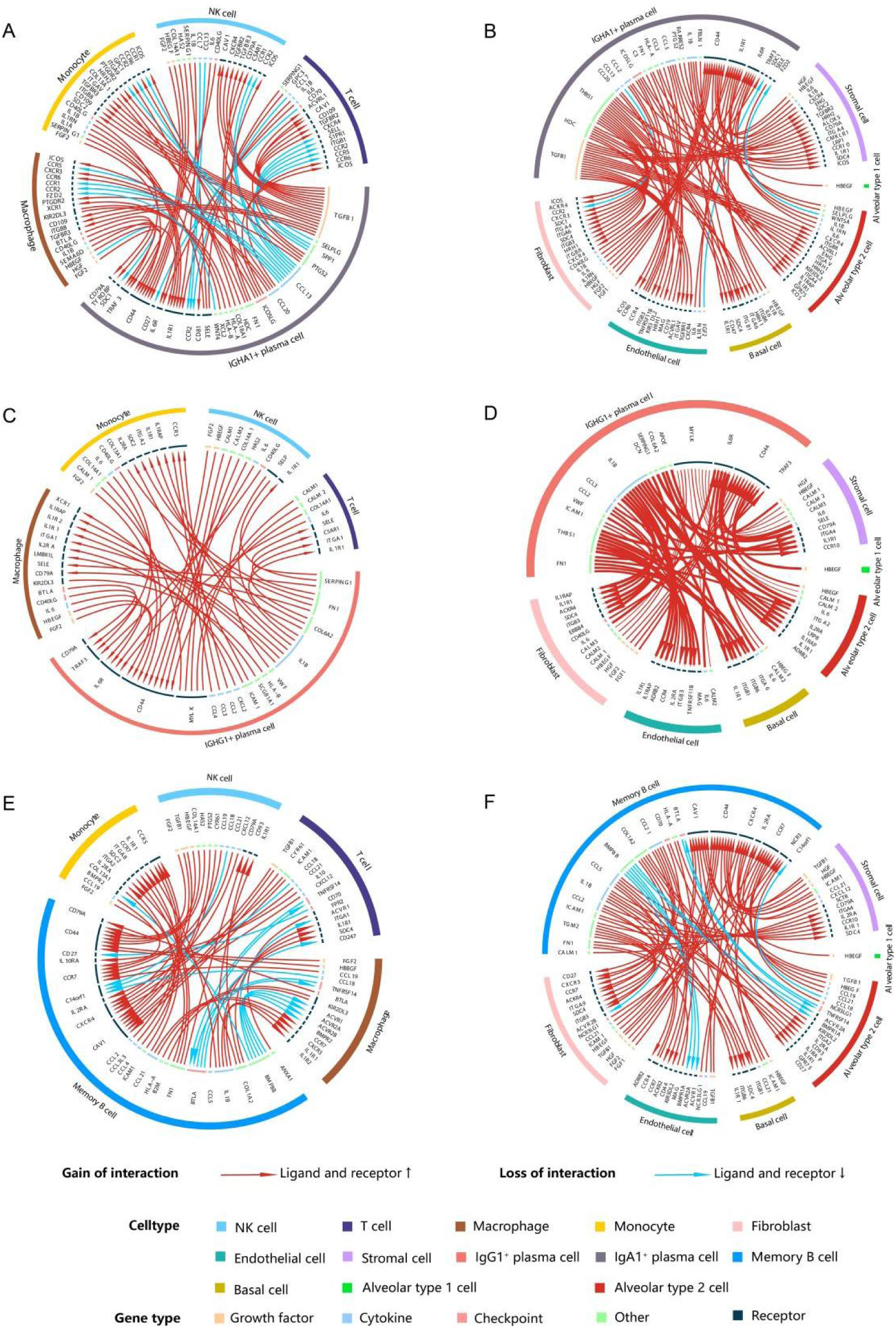
Cell–cell communication between B-cell subsets and the pulmonary microenvironment in COPD. Cell–cell communication analyses were performed to investigate predicted interactions between IgG1^+^ plasma cells (A–B), IgA1^+^ plasma cells (C–D), and memory B cells (E–F) and surrounding immune-cell populations, including T cells, monocytes, macrophages, and NK cells, as well as non-immune cell populations, including fibroblasts, endothelial cells, stromal cells, basal cells, and alveolar epithelial cells, in the lungs of patients with COPD.

Similarly, IgG1^+^ plasma cells demonstrated interactions involving IL-1B/IL-6 signalling, collagen–integrin pathways and HLA-B-associated communication with monocytes and macrophages. Enhanced chemokine signalling (CCL2/3/4–CCR5) was associated with monocyte recruitment. Signals derived from neighbouring immune-cell populations were associated with increased CD44, MYLK, IL6R, and TRAF3 expression on IgG1^+^ plasma cells (Figure 3C–D; Supplementary Table 3).

Memory B cells also exhibited extensive interactions with surrounding immune and stromal cells, predominantly through IL-1B-related signalling pathways. In addition, predicted FN1- and CCL21-associated signalling suggested enhanced communication with monocytes and macrophages. Reciprocal interactions from immune cells were associated with increased CAV1, CCR7, and CD44 expression in memory B cells (Figure 3E–F, Supplementary Table 3). Collectively, these findings suggest that B-cell subsets in COPD may contribute to persistent inflammatory activation and tissue remodelling within the lung microenvironment.

### 3.3 Transitional B cells link peripheral immune dysregulation with COPD-associated autoimmunity

To explore systemic B-cell abnormalities in COPD, we analysed peripheral blood B cells (GSE205078), classifying them into seven major subclusters [21,27] (Figure 4A–D, Supplementary Figure 2). Compared with controls, COPD samples exhibited increased proportions of naïve B cells and reduced proportions of transitional B cells (Figure 4B).

**Figure 4.**
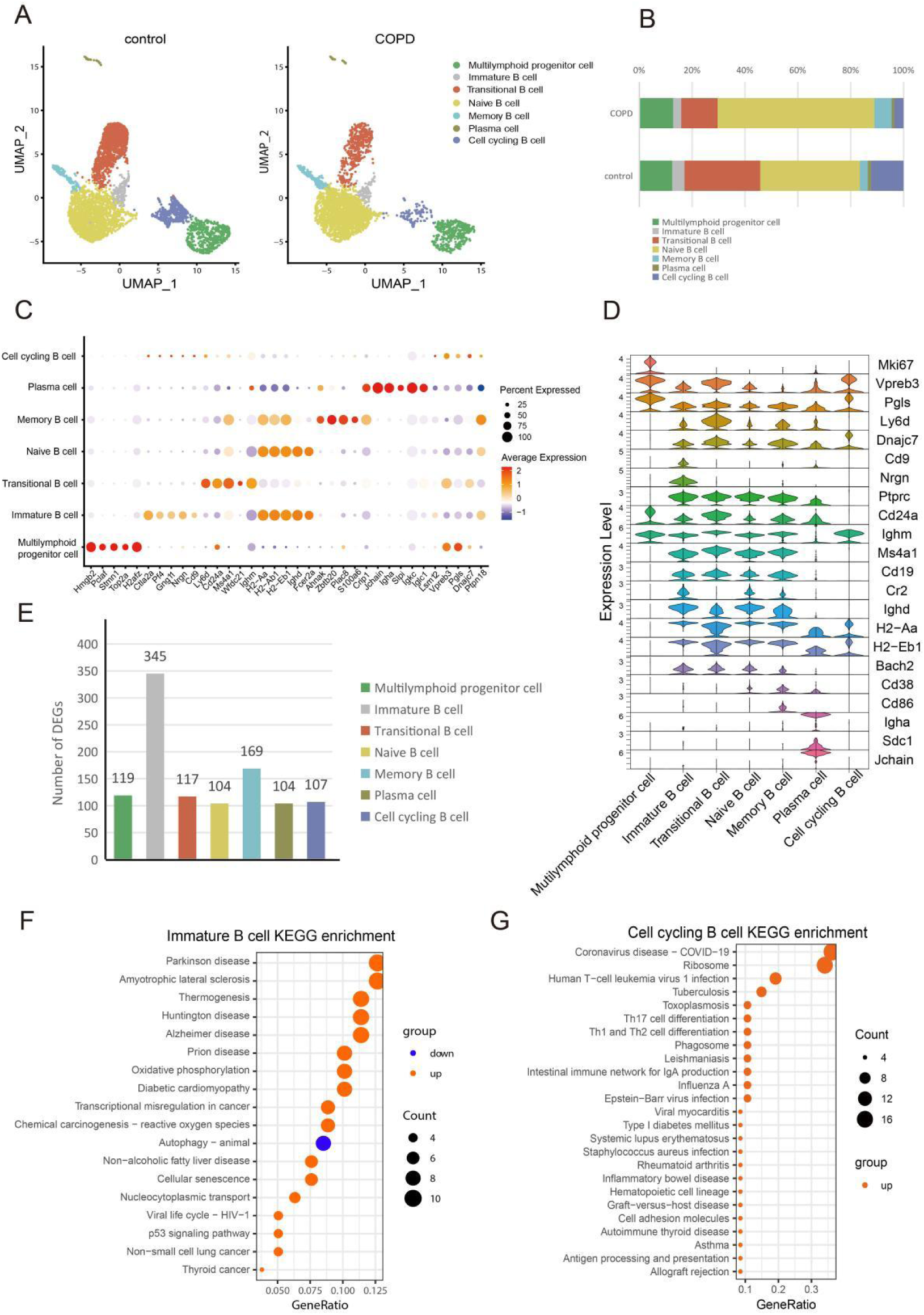
Peripheral blood B-cell subclusters in COPD mice. **(A)** Uniform Manifold Approximation and Projection (UMAP) visualization of peripheral blood B cells derived from COPD (n = 4,400) and control (n = 3,228) mice, coloured according to B-cell subsets. **(B)** Relative frequencies of individual B-cell subsets in COPD and control mice. **(C)** Dot plot showing the top five marker genes for each B-cell subset. **(D)** Violin plots displaying representative marker-gene expression across B-cell subsets. **(E)** Bar plot showing the numbers and proportions of differentially expressed genes (DEGs) identified in each B-cell subset. **(F–G)** Kyoto Encyclopedia of Genes and Genomes (KEGG) pathway enrichment analyses of upregulated (red) and downregulated (blue) pathways in immature B cells (F) and cell-cycling B cells (G). Circle size represents the number of genes enriched within each pathway.

Differential gene expression analysis revealed significant transcriptional alterations in immature and memory B-cell populations (Figure 4E). KEGG analysis demonstrated enrichment of pathways associated with immune activation and inflammatory responses in immature B cells (Figure 4F). Cell-cycling B cells exhibited enrichment of autoimmune-associated pathways, including SLE, RA, autoimmune thyroid disease (AITD), IBD, Th1/Th2/Th17 differentiation, B-cell activation, and intestinal IgA production pathways (Figure 4G). In contrast, autoimmune-associated pathways were not prominently enriched in other B-cell subsets (Supplementary Figure 3A–C).

Because transitional B cells represent a critical developmental stage linking immature bone marrow B cells to mature peripheral B cells and are frequently altered in autoimmune diseases [28], these populations were further deeply analysed. PPI network analysis showed predominantly downregulated proteins, including Uba52, Rpl29, and Gna13 (Figure 5A). KEGG analysis confirmed enrichment of multiple autoimmune-associated pathways, including RA, T1DM, SLE, IBD, AITD, and Th17 differentiation pathways (Figure 5B).

**Figure 5.**
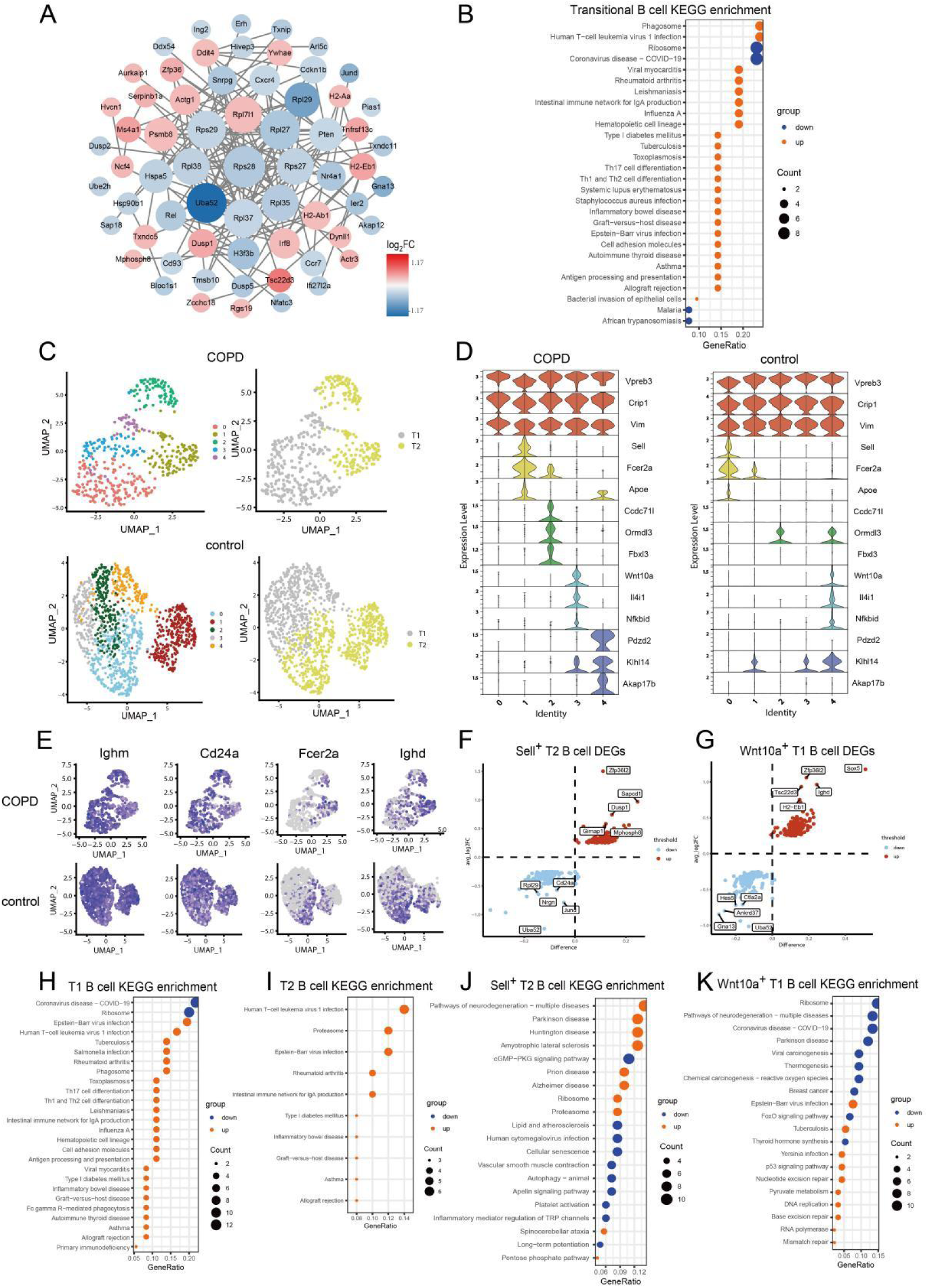
Transitional B-cell abnormalities in the peripheral blood of COPD mice. **(A)** Protein–protein interaction (PPI) network of differentially expressed genes (DEGs; |log₂FC| >0.6) identified in transitional B cells. Red nodes indicate upregulated genes, whereas blue nodes indicate downregulated genes. **(B)** Kyoto Encyclopedia of Genes and Genomes (KEGG) pathway enrichment analysis of upregulated (red) and downregulated (blue) pathways in transitional B cells. Circle size represents the number of genes enriched within each pathway. **(C)** Uniform Manifold Approximation and Projection (UMAP) visualization of transitional B cells from COPD and control mice coloured according to cluster identity and developmental subsets. **(D)** Violin plots showing representative marker-gene expression across transitional B-cell subclusters. **(E)** Expression patterns of transitional B-cell classification markers, including Ighm, Cd24a, Fcer2a, and Ighd. **(F–G)** Volcano plots of differentially expressed genes in Sell^+^ T2 B cells (F) and Wnt10a^+^ T1 B cells (G). Red dots indicate upregulated genes, whereas blue dots indicate downregulated genes. **(H–K)** KEGG pathway enrichment analyses of upregulated (red) and downregulated (blue) pathways in T1 B cells (H), T2 B cells (I), Sell^+^ T2 B cells (J), and Wnt10a^+^ T1 B cells (K). Circle size indicates the number of enriched genes within each pathway.

Transitional B cells in COPD mice were further divided into T1 and T2 subsets based on *Ighm*, *Ighd*, *Cd24a*, and *Fcer2a* (*Cd23*) expression patterns [25,27] (Figure 5C–E). COPD mice exhibited expansion of T1 transitional B cells (Figure 5C, E). Furthermore, clusters in COPD mice, characterised by Wnt10a expression (Wnt10a^+^ T1) and Sell expression (Sell^+^ T2), corresponded to distinct transitional-cell populations identified in control animals (Figure 5D, F, G). Both T1 and T2 transitional B cells exhibited enrichment of autoimmune-associated pathways, including RA, T1DM, and IBD (Figure 5H–I). In addition, Sell^+^ T2 and Wnt10a^+^ T1 B cells exhibited enrichment of pathways associated with oxidative stress, mitochondrial dysfunction, DNA damage responses, macrophage activation and inflammatory signalling, whereas FoxO signalling pathways were relatively suppressed in Wnt10a^+^ T1 B cells (Figure 5J–K). Similar to the findings in mice, T1 B cells from COPD patients were prominently enriched in autoimmune disease related pathways, including RA, IBD, autoimmune thyroid disease (AITD), T1DM and graft-versus-host disease. Th1/Th2 as well as Th17 cell differentiation pathways and those of the intestinal immune network for IgA production were also enriched, suggesting enhanced immune activation and inflammatory responsiveness in T1 B cells in COPD patients (Table 2, Supplementary Figure 4). Taken together, these findings suggest that transitional B-cell dysregulation may represent an important link between peripheral immune abnormalities and COPD-associated autoimmunity.

### 3.4 Early B-cell development is altered in the bone marrow of COPD mice

Given that receptor editing and negative selection occur during early B-cell development in the bone marrow, we next investigated whether central B-cell tolerance was altered in COPD. Bone marrow B cells of COPD mice (GSE205078) were classified into seven subsets (Supplementary Figure 5A, C, E; Figure 6A–D). Compared with controls, COPD mice showed reduced proportions of small pre-B III cells and increased proportions of pre-B I, immature B, and mature B cells (Figure 6B).

**Figure 6.**
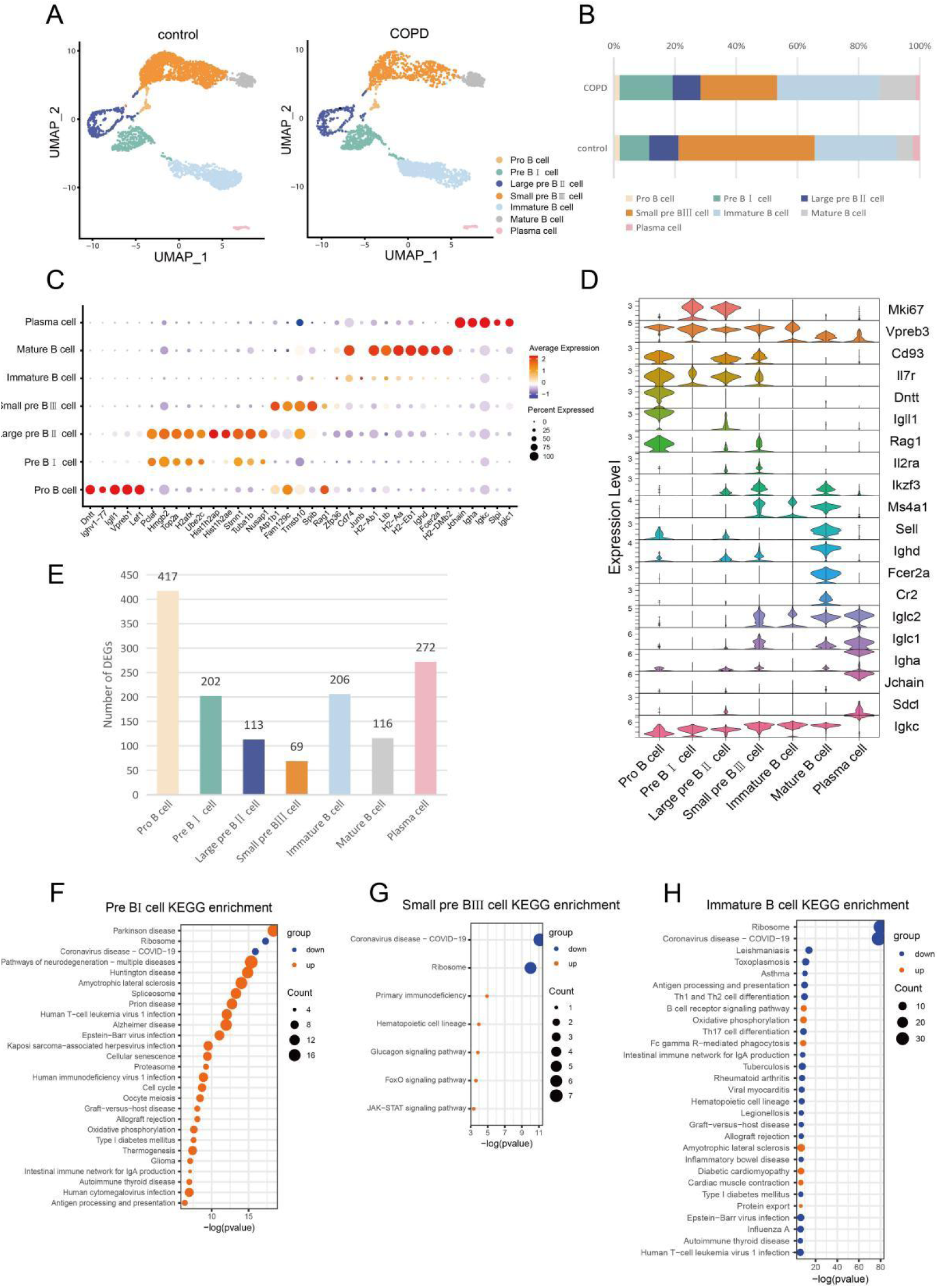
Bone marrow B-cell subclusters in COPD mice. **(A–B)** Uniform Manifold Approximation and Projection (UMAP) visualization of bone marrow B cells from control (n = 3,707) and COPD (n = 2,816) mice, together with the relative proportions of individual B-cell subsets. Cells are coloured according to subset identity. **(C)** Dot plot showing the top five marker genes for each bone marrow B-cell subset. **(D)** Violin plots displaying representative functional marker-gene expression across B-cell subsets. **(E)** Bar plot showing the numbers of differentially expressed genes (DEGs) identified in each B-cell subset. **(F–H)** Kyoto Encyclopedia of Genes and Genomes (KEGG) pathway enrichment analyses of upregulated (red) and downregulated (green) pathways in pre-B I cells (F), small pre-B III cells (G), and immature B cells (H). Circle size represents the number of genes enriched within each pathway.

DEGs in each subset were showed in Figure 6E. KEGG analysis of DEGs revealed that pre-B I cells were enriched for autoimmune-associated pathways, including T1DM, AITD, graft-versus-host disease and intestinal IgA network pathways (Figure 6F). In contrast, immature B and large pre-B II cells exhibited downregulation of several autoimmune-associated pathways despite enhanced BCR signalling-related pathways (Figure 6H; Supplementary Figure 3D). Small pre-B III cells showed enrichment of pathways associated with B-cell differentiation and development, including FoxO and JAK–STAT signalling pathways (Figure 6G). These findings are compatible with the hypothesis that early B-cell developmental processes and central tolerance checkpoints are altered in COPD.

### 3.5 COPD is associated with altered BCR repertoire remodelling and λ-chain enrichment

To further investigate BCR abnormalities associated with COPD, we performed single-cell V(D)J sequencing on human peripheral blood (Table 2) and identified 9 major B-cell subsets (Figure 7A–C). Compared with controls, COPD samples exhibited reduced proportions of resting naïve (rNAV) and resting memory (rM) B cells together with increased proportions of transitional (Tran), early activated naïve (eaNAV), and activated class-swiched memory (SA-MEM) B cells (Figure 7B–C).

**Figure 7.**
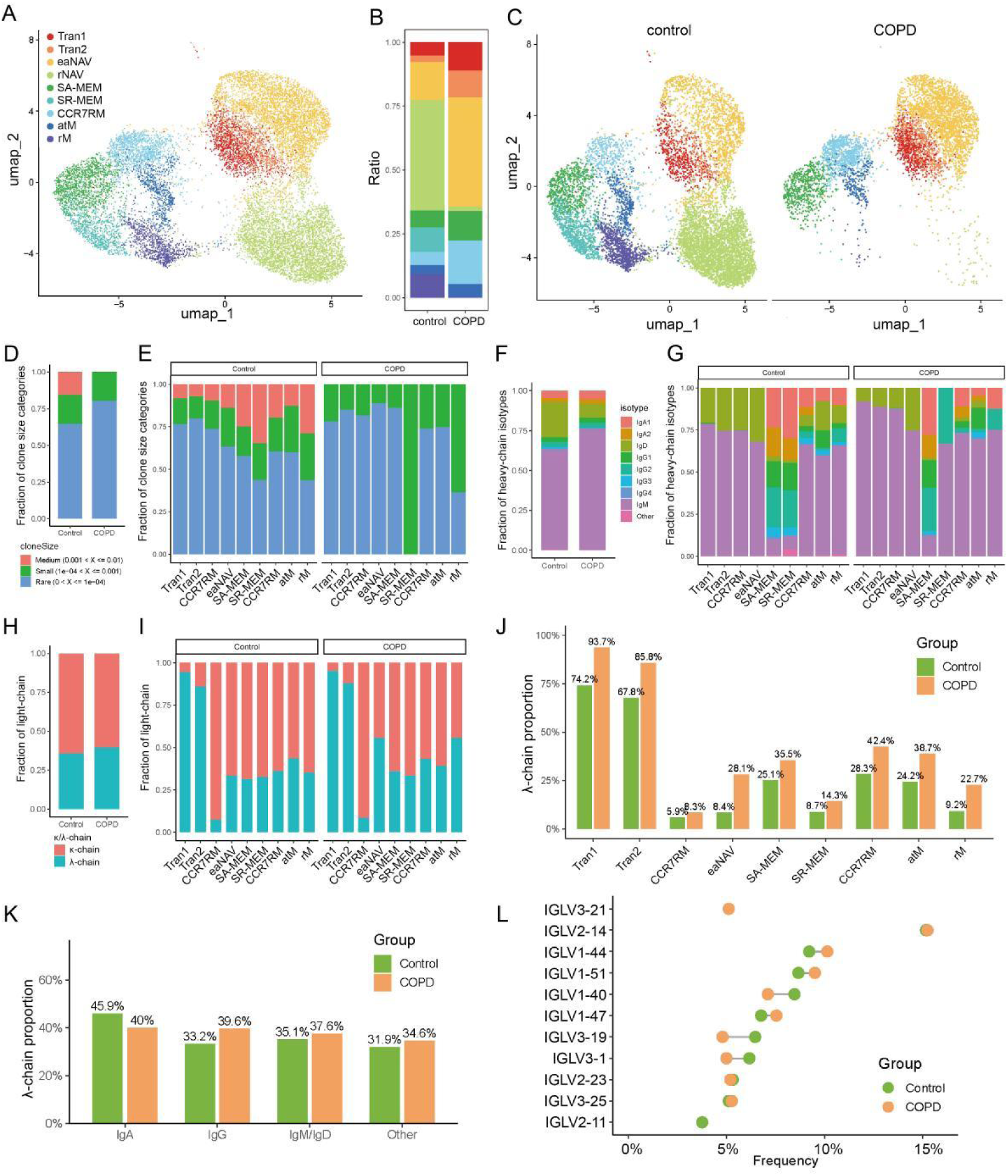
BCR repertoire remodelling and altered light-chain usage in patients with COPD. **(A–C)** Uniform Manifold Approximation and Projection (UMAP) visualization (A, C) and relative proportions (B) of peripheral blood B cells from healthy controls (n = 9,942) and patients with COPD (n = 5,474), coloured according to annotated B-cell subsets. **(D–E)** Distribution (D) and proportions (E) of B-cell clonotype sizes in control and COPD samples. **(F–G)** Immunoglobulin isotype distributions shown overall (F) and across individual B-cell subsets (G). **(H–I)** κ/λ light-chain distributions shown overall (H) and within individual B-cell subsets (I). **(J–K)** Relative proportions of λ light-chain usage across B-cell subsets (J) and immunoglobulin isotypes (K). **(L)** Top 10 IGLV gene usages identified in control and COPD samples. Abbreviation: Tran, Transitional; eaNAV, early activated naïve; rNAV, resting naïve; SA-MEM, activated class-swiched memory; SR-MEM, resting class-swiched memory; RM, recirculing memory; atypical memory, atM; rM, resting memory.

BCR repertoire analysis revealed an increased proportion of rare clonotypes and a reduced proportion of medium-sized clonotypes in COPD patients. Resting class-switched memory (SR-MEM) B cells were composed predominantly of small clonotypes (Figure 7D–E). Isotype analysis further revealed increased IgM and reduced IgD proportions in COPD samples compared with controls (Figure 7F–G).

Light-chain analysis demonstrated increased λ-chain usage in COPD without evidence of dominant monoclonal expansion. Stacked bar plots further revealed a global shift towards λ-chain usage across multiple B-cell subsets in COPD. Notably, λ chains were predominantly enriched in transitional B cells, whereas κ chains were more highly represented in naïve and memory B-cell populations (Figure 7H–I). These findings suggest altered B-cell selection and BCR repertoire remodelling in COPD, consistent with the transitional B-cell abnormalities identified by scRNA-seq analyses (Figures 4–5; Supplemeantary Figure 4).

In addition, increased IgG isotypes and reduced IgA isotypes were preferentially associated with λ-chain expression across multiple B-cell subsets in COPD (Figure 7J–K). These findings indicate altered relationships between immunoglobulin isotypes, light-chain usage, and B-cell subsets in COPD.

Analysis of the top 10 IGLV genes demonstrated enrichment of several λ-chain V genes, including IGLV1-47, in COPD samples (Figure 7L), further supporting light-chain repertoire remodelling under chronic inflammatory conditions. Global diversity analysis additionally demonstrated increased BCR diversity in COPD B cells. Memory B cells exhibited elevated diversity indices (Shannon entropy and inverse Simpson index) together with increased richness estimates (Chao1 and ACE), suggesting broad polyclonal activation rather than dominant oligoclonal expansion (Supplementary Figure 6).

To further investigate whether altered BCR development contributes to these abnormalities, v-Abl/Bcl2 pro-B cells were stimulated with CSE *in vitro*. CSE treatment significantly reduced λ5 expression in a concentration-dependent manner (Figure 8B), supporting the impairment of early B-cell development in COPD.

**Figure 8.**
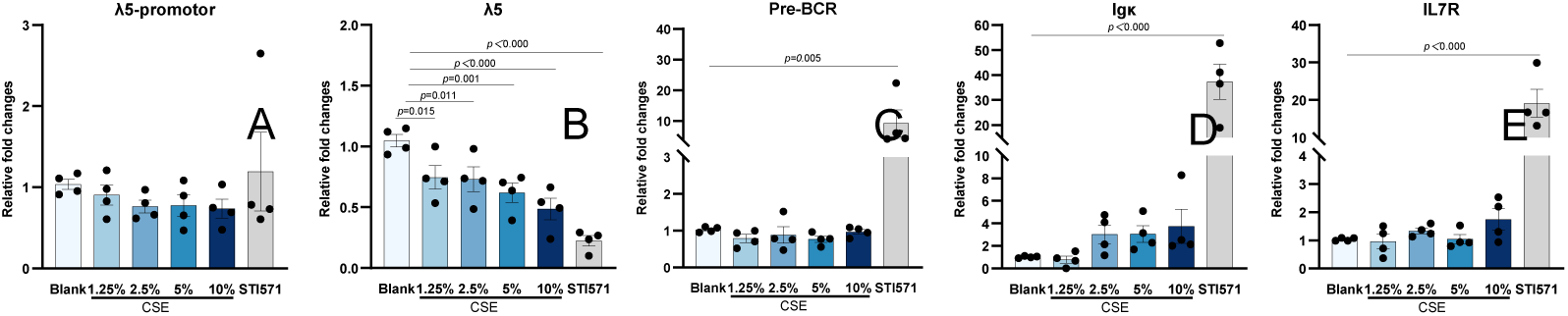
Cigarette smoke extract suppresses λ5 expression in a v-Abl/Bcl2 pro-B cell line. V-Abl/Bcl2 pro-B cells were treated with cigarette smoke extract (CSE; 1.25–10%) for 48 h. STI571 (3 μM) was used as a positive control. Reverse transcription PCR (RT–PCR) was performed to measure the expression of λ5-promoter, λ5, κ, pre-BCR, and IL-7R mRNA. β-actin served as the internal loading control. Data are presented as mean ± standard error of the mean (SEM) (n = 4). p values were calculated relative to the untreated control group.

### 3.6 COPD exhibits restricted and low-grade autoimmune features compared with classical autoimmune diseases

To contextualise B-cell abnormalities in COPD, comparative analyses were performed using datasets from SSc lung tissue (GSE132771), SLE (GSE193867) peripheral blood and RA bone marrow samples (GSE221704, Supplementary Figures 1, 2, 5, and 7–11).

Comparison of lung tissues between COPD and SSc indicated that IgA1^+^ plasma cells were reduced in SSc, whereas IgG1^+^ plasma cells were increased and memory B cells were reduced in SSc, contrasting with the trends observed in COPD (Figure 1D; Figure 9A). Shared transcriptional features included genes associated with antigen presentation (*HLA-B*) and B-cell dysregulation (*Ddnt* and *Zfp* family genes). *IGHG4* expression was commonly upregulated in memory B cells across both diseases (Figure 9D–F, Supplementary Table 4). KEGG analyses demonstrated overlapping enrichment of macrophage activation and autoimmune-associated pathways, including T1DM and graft-versus-host disease pathways (Figure 1I; Supplementary Figure 7H). Cell–cell communication analysis revealed stronger HLA-associated signalling in SSc, whereas COPD demonstrated relatively greater enrichment of inflammation- and remodelling-associated signalling pathways (Figure 3; Supplementary Tables 3 and 5; Supplementary Figure 8).

**Figure 9.**
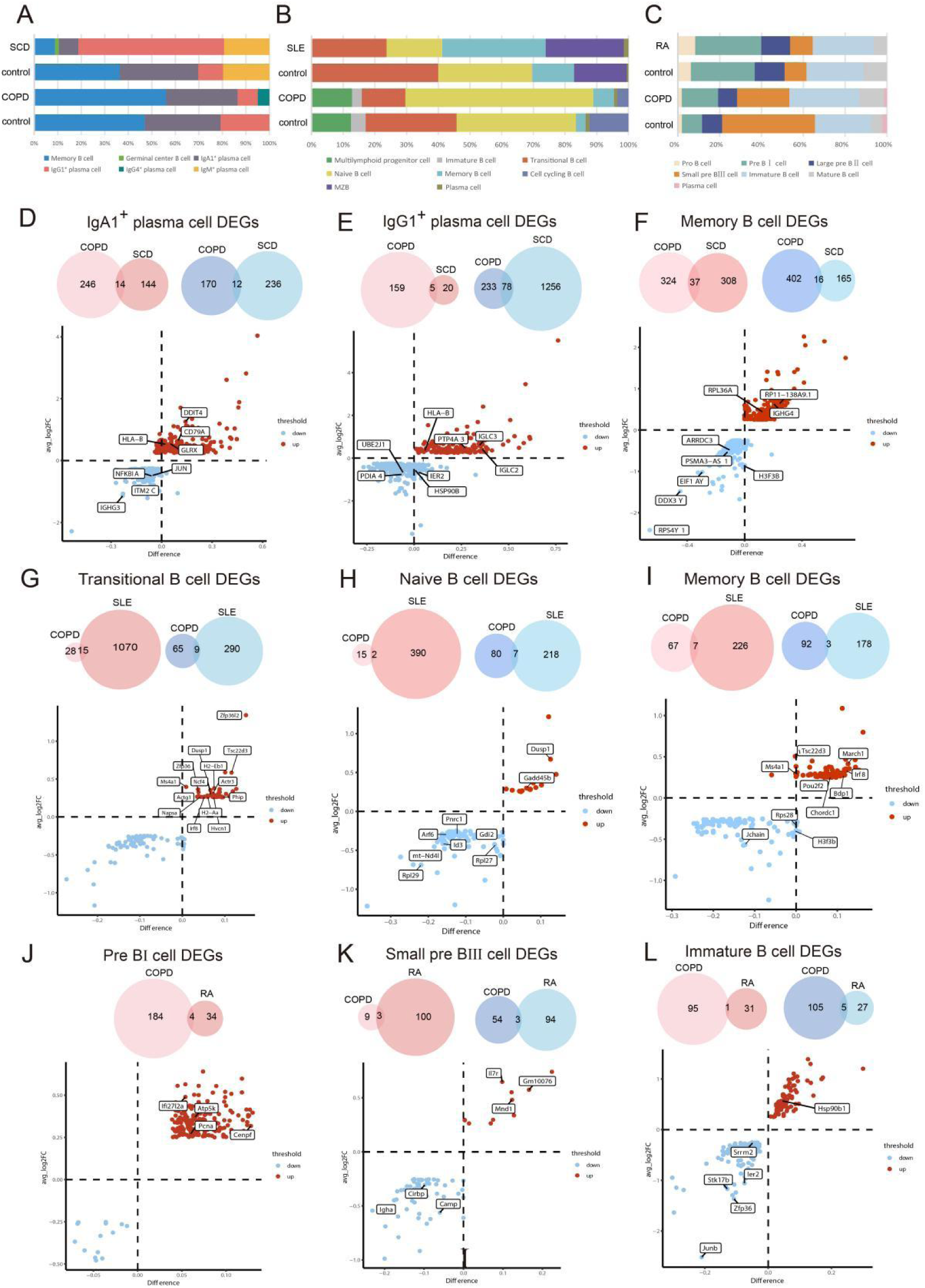
Comparative analysis of B-cell autoimmune abnormalities in COPD and classical autoimmune diseases. **(A–C)** Relative proportions of B-cell subsets in the lungs of patients with COPD and systemic sclerosis (SSc) (A), in the peripheral blood of COPD mice and patients with systemic lupus erythematosus (SLE) (B), and in the bone marrow of COPD and rheumatoid arthritis (RA) mice (C). **(D–L)** Volcano plots showing differentially expressed genes (DEGs) in IgA1^+^ plasma cells (D), IgG1^+^ plasma cells (E), and memory B cells (F) from the lungs of patients with COPD and SSc; transitional B cells (G), naïve B cells (H), and memory B cells (I) from the peripheral blood of COPD mice and patients with SLE; and pre-B I cells (J), small pre-B III cells (K), and immature B cells (L) from the bone marrow of COPD and RA mice. Shared DEGs between COPD and the corresponding autoimmune diseases are highlighted. Venn diagrams indicate the numbers of total and overlapping DEGs within each subset. Upregulated and downregulated genes are shown in red and blue, respectively.

Comparison of peripheral blood samples between COPD mice and patients with SLE revealed reduced transitional B-cell proportions together with increased memory B-cell proportions (Figure 4B; Figure 9B; Supplementary Figure 9B). KEGG analyses demonstrated overlapping enrichment of autoimmune-associated pathways, antigen presentation pathways and Th1/Th2 differentiation pathways within transitional B-cell populations (Figure 5H–I; Supplementary Figures 9 and 10). Shared differentially expressed genes included *H2-Eb1, H2-Aa,* and *Zfp36* in transitional B cells; *Dusp1* in transitional and naïve B cells; and *Irf8* in transitional and memory B cells (Figure 9G–I, Supplementary Table 4).

In bone marrow, comparisons between COPD mice and collagen-induced arthritis (CIA) mice showed that the proportions of pre-B I, pre-B III and immature B cells were relatively preserved in CIA mice compared with controls, in contrast to the marked developmental alterations observed in COPD mice (Figure 6B; Figure 9C; Supplementary Figure 11B). In RA mice, autoimmune-associated pathway enrichment was primarily observed in pre-B III and immature B-cell populations, whereas in COPD mice enrichment was predominantly observed in pre-B I cells (Figure 9J–L; Supplementary Figure 11F–G). Shared upregulated genes included *Ifi27l2a, Pcna, Atp5k,* and *Cenpf* in pre-B I cells; *Il7r, Mnd1,* and *Gm10076* in small pre-B III cells; and *Hsp90b1* in immature B cells (Figure 9J–L, Supplementary Table 4).

Collectively, these comparative analyses suggest that COPD shares several immunological features with classical autoimmune diseases but manifests a more restricted and low-grade autoimmune phenotype characterised by chronic inflammatory activation, polyclonal B-cell responses and local tissue remodelling rather than systemic autoimmunity.

## 4 Discussion

Since the autoimmune hypothesis of COPD was first proposed over two decades ago [3], accumulating evidence has supported the involvement of autoimmune dysregulation in COPD pathogenesis. The mechanisms linking chronic inflammation, B-cell abnormalities and tissue remodelling however remain incompletely understood. In the present study, through integration of single-cell transcriptomic analysis, single-cell BCR sequencing and *in vitro* validation experiments, we identified a COPD-associated B-cell immune signature characterised by enrichment of IgG4^+^ plasma cells, altered transitional B-cell populations, aberrant bone marrow B-cell development and λ-chain-biased BCR remodelling. Collectively, these findings suggest that persistent low-grade B-cell autoimmunity may contribute to chronic inflammatory injury and remodelling in COPD.

A major finding of this study was the identification of a distinct, selectively enriched IgG4^+^ plasma-cell population in COPD lung tissue associated with increased expression of *IGHG4* together with genes related to BCR activation and inflammatory-response pathways (Figure 1). IgG4 antibodies are classically associated with IgG4-related disease (IgG4-RD) and IgG4-mediated autoimmune disorders [29]. In IgG4-RD, the expansion of IgG4^+^ plasma cells contributes to fibro-inflammatory tissue injury and chronic immune activation [30]. Unlike other IgG subclasses, IgG4 antibodies can undergo Fab-arm exchange, generating functionally bispecific antibodies capable of recognising diverse antigens. Although the precise role of IgG4 in COPD remains unclear, the enrichment of IgG4^+^ plasma cells, coupled with the increased serum IgG4 concentrations and their inverse trends with pulmonary function parameters is compatible with the hypothesis that IgG4-associated immune activation contributes to local inflammatory injury in COPD lungs.

Of the highly expressed genes in IgG4^+^ plasma cells, *IGLV1-47* emerged as a notable member. This λ-chain V-gene segment was also enriched in peripheral blood BCR sequencing data, suggesting coordinated light-chain repertoire remodelling across pulmonary and systemic compartments. Although the biological function of *IGLV1-47* remains poorly characterised, previous studies have linked this gene to severe asthma associated with gastroesophageal reflux disease [31]. The enrichment of *IGLV1-47* in COPD therefore raises the possibility that chronic inflammatory stimulation and altered antigen selection contribute to λ-chain-biased B-cell activation. Furthermore, increased BCL2 expression, together with enrichment of TNF, MAPK, and NF-κB signalling pathways, supports the presence of persistent inflammatory activation and enhanced survival signalling within COPD-associated B-cell populations. Nevertheless, the pathogenic role of IgG4 antibodies and *IGLV1-47*-enriched B cells in COPD requires direct functional validation.

Another important finding was the consistent upregulation of *MT2A* across multiple pulmonary B-cell subsets, including IgA1^+^ plasma cells, IgG1^+^ plasma cells and memory B cells. Interestingly, our previous work identified a population of MT-high T cells in COPD lungs and peripheral blood characterised by elevated metallothionein expression, including *MT2A* [33]. In that study, MT-high T cells exhibited immunoregulatory properties and progressively declined during COPD progression, suggesting a protective role in limiting excessive inflammation. In contrast, the present study demonstrates increased *MT2A* expression in pulmonary B-cell populations. These divergent findings likely reflect cell-type-specific responses to chronic oxidative stress and inflammatory signalling within the COPD microenvironment.

Metallothioneins are important regulators of zinc homeostasis, oxidative stress responses and inflammatory signalling [34,35]. In RA, *MT2A* promotes inflammatory metabolic reprogramming and IL-1β production in synovial monocytes [35]. Consistent with these observations, increased *MT2A* expression in COPD B cells, along with enrichment of inflammatory chemokine pathways including CCL3 and CCL4 signalling, suggests that *MT2A* may participate in monocyte/macrophage recruitment and chronic inflammatory remodelling in COPD lungs. Nevertheless, because the present study is based primarily on transcriptomic and BCR repertoire analyses, the functional role of *MT2A* in COPD B cells remains speculative and requires further mechanistic validation through dedicated *in vitro* and *in vivo* studies.

Importantly, transitional B-cell abnormalities were consistently observed across both murine COPD surrogates and human peripheral blood, suggesting that these findings are unlikely to represent species-specific phenomena. The expansion of T1 transitional B cells together with altered expression of Wnt10a and Sell, genes previously implicated in autoimmune disease pathogenesis [36], supports the concept that peripheral B-cell maturation is altered in COPD. The enrichment of autoimmune-associated pathways within transitional B-cell populations further suggests that these cells may reflect an intermediate immune state linking central B-cell developmental abnormalities with peripheral immune dysregulation.

Bone marrow analyses provided additional evidence linking the pathogenesis of COPD with abnormalities in early B-cell development. The bone marrow of the murine COPD surrogates demonstrated expansion of pre-B I cells, reduction of small pre-B III cells and altered expression of genes involved in receptor editing and B-cell maturation, including *Il7r, Vpreb1/2*, and *Rag* family genes. Because receptor editing and negative selection are critical mechanisms for maintaining central B-cell tolerance, these findings raise the possibility that self-reactive B cells may escape early developmental checkpoints and subsequently contribute to peripheral autoimmune activation in COPD. This interpretation is further supported by the observed λ-chain enrichment and altered BCR repertoire characteristics identified in peripheral blood B cells.

Our BCR sequencing analyses demonstrated that COPD is characterised by increased λ-chain usage, enrichment of *IGLV1-47*, an altered immunoglobulin isotype distribution and increased repertoire diversity without evidence of dominant monoclonal expansion. These findings suggest broad, polyclonal B-cell activation rather than classical oligoclonal autoimmune responses. In particular, λ-chain enrichment was preferentially associated with transitional B cells and activated B-cell populations, whereas κ chains remained more abundant within naïve and resting memory compartments. Together, these findings support the possibility that altered receptor editing and chronic antigenic stimulation contribute to abnormal B-cell selection in COPD.

Previous studies using porcine pancreatic elastase (PPE)-induced murine surrogates of emphysema demonstrated that B1 cells, particularly the B1b subset, contribute to alveolar destruction through ADAM10-mediated mechanisms [37]. Our findings extend these observations by demonstrating that COPD-associated B-cell abnormalities are not restricted to innate-like B1 populations but also involve conventional B2-cell subsets, including plasma cells, memory B cells, and transitional B cells across multiple immune compartments.

Comparative analyses with SSc, SLE, and RA further revealed that COPD shares several immunological characteristics with classical autoimmune diseases while retaining distinct features. Shared transcriptional signatures included HLA-associated antigen presentation pathways, *IGHG4* upregulation, interferon-related inflammatory responses and transitional B-cell abnormalities. Several important differences did, however, emerge: in SLE, autoimmune-associated pathways were broadly enriched across multiple B-cell populations, whereas in COPD these abnormalities appeared more spatially and developmentally restricted. Similarly, the transcriptional signatures of the B cells implicated in COPD pathogenesis exhibited wider repertoire diversity and polyclonal activation in the absence of dominant monoclonal expansion, contrasting with the more pronounced autoreactive clonal responses frequently observed in classical autoimmune diseases. In the bone marrow, the B cell signatures in COPD and RA shared evidence of altered receptor-editing-related pathways, yet the affected developmental stages and associated transcriptional programmes differed substantially. These observations suggest that COPD may reflect the outcome of a distinct form of chronic, low-grade autoimmunity characterised by persistent inflammatory activation and local tissue remodelling rather than overt systemic autoimmune disease.

Several limitations should be acknowledged. Firstly, integrated single-cell datasets simultaneously covering lung tissue, peripheral blood, and bone marrow from both human subjects and matched animal models remain limited. Secondly, the number of autoimmune diseases included for comparative analysis in our study was relatively small, precluding validation of several of our observations in independent patient cohorts. Thirdly, although our findings identify strong associations between B-cell abnormalities and COPD pathogenesis, direct causal relationships between specific B-cell subsets and disease progression remain to be established. In particular, the precise pathogenic roles of IgG4^+^ plasma cells, λ-chain-biased BCR remodelling, and *MT2A* upregulation require functional investigation.

In conclusion, this study provides multi-dimensional evidence linking IgG4^+^ plasma-cell enrichment, λ-chain-biased BCR remodelling and altered transitional B-cell development to COPD-associated autoimmunity. Our findings support a model in which abnormalities in early B-cell development and receptor editing contribute to persistent, low-grade autoimmune activation in COPD. Compared with classical autoimmune diseases, COPD appears to exhibit a more restricted and predominantly localised autoimmune phenotype, characterised by chronic inflammatory remodelling and broad polyclonal B-cell activation. Together, these findings expand the current understanding of B-cell-mediated immune dysregulation in COPD and provide a potential framework for future immune-targeted therapeutic strategies.

## Supporting information

IgG4 and BCR remodeling in COPD

## Data Availability

The single-cell BCR sequencing data generated in this study have been deposited in the Genome Sequence Archive (GSA) of the National Genomics Data Center (https://ngdc.cncb.ac.cn/gsa-human) under accession number HRA018181 (Project ID: PRJCA063022). These data will be released upon publication. All other data supporting the findings of this study are available within the article and its supplementary information files, or from the corresponding authors upon reasonable request.

## Competing Interests

The authors declare no competing interests.

## Funding

This work was supported by the Beijing Natural Science Foundation (grant no. 7242002) and the National Natural Science Foundation of China (grant no. 82090013) and sponsored by Beijing Nova Program (No. 20240484639). The funders had no role in the study design, data collection, data analysis, data interpretation, manuscript preparation or the decision to submit the work for publication.

## Acknowledgement

We thank Professor Jiazhi Hu from the School of Life Sciences, Peking University, for generously providing the v-Abl/Bcl2 pro-B cell line used in this study.

## Author Contributions (CRediT)

H.H.Y. conceived the research project following initial discussions with W.W., Y.C. and Y.S. L.D. led the single-cell RNA-seq analyses. H.Z. and Y.Q.X. contributed to early single-cell data analysis and interpretation. Z.Q.L. and M.M. collected human samples. L.D. and H.H.Y. performed the *in vitro* experiments. H.H.Y. drafted the initial version of the manuscript with assistance from H.Z. and Y.Q.X. Y.S., and W.W. reviewed the data and critically revised the manuscript. C. J. Corrigan offered valuable comments on the manuscript. All authors read and approved the final version of the manuscript.

## Ethics Approval

Human specimen collection was approved by the Institutional Review Board of Capital Medical University (approval no. Z2021SY025). Written informed consent was obtained from all participants prior to enrolment.

## Notes

### Competing Interest Statement

The authors have declared no competing interest.

