## Supplementary material for "IgG4^+^ plasma cell enrichment and λ-chain-biased BCR remodeling drive low-grade autoimmunity in chronic obstructive pulmonary disease": IgG4 and BCR remodeling in COPD

**Supplementary Figures and legends**

**Supplementary Figure 1**

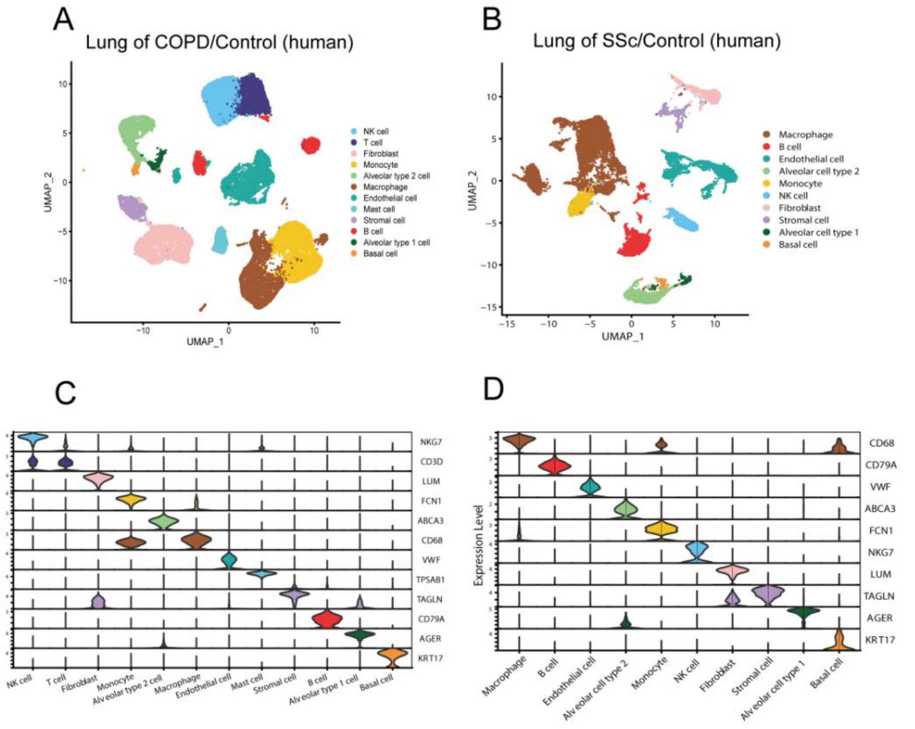


E

| cluster Name | Cell number | | | | % of total | | | |
|---|---|---|---|---|---|---|---|---|
| | Control | COPD | Control | SCD | Control | COPD | Control | SCD |
| B cell | 483 | 1324 | 66 | 1452 | 3.52% | 6.72% | 0.88% | 19.71% |
| Macrophage | 2541 | 1346 | 1997 | 4052 | 18.50% | 6.83% | 26.66% | 55.01% |
| Monocyte | 2381 | 2743 | 874 | 30 | 17.34% | 13.93% | 11.67% | 0.41% |
| NK cell | 1734 | 4416 | 526 | 221 | 12.62% | 22.42% | 7.02% | 3.00% |
| T cell | 1541 | 2621 | 0 | 0 | 11.22% | 13.31% | 0.00% | 0.00% |
| Endothelial cell | 1843 | 2331 | 1517 | 981 | 13.42% | 11.84% | 20.25% | 13.32% |
| Fibroblast | 1354 | 2643 | 842 | 213 | 9.86% | 13.42% | 11.24% | 2.89% |
| Stromal cell | 295 | 934 | 340 | 242 | 2.15% | 4.74% | 4.54% | 3.29% |
| Basal cell | 142 | 58 | 38 | 103 | 1.03% | 0.29% | 0.51% | 1.40% |
| Alveolar cell type 1 | 44 | 447 | 286 | 1 | 0.32% | 2.27% | 3.82% | 0.01% |
| Alveolar cell type 2 | 1377 | 832 | 1006 | 71 | 10.03% | 4.22% | 13.43% | 0.96% |
| total | 13735 | 19695 | 7492 | 7366 | | | | |

**Supplementary Figure 1. Single-cell transcriptomic landscape of lung tissues from patients with COPD and systemic sclerosis (SSc). (A–B)** Integrated Uniform Manifold Approximation and Projection (UMAP) visualization of lung cells derived from patients with COPD (n = 33,430 cells) and systemic sclerosis (SSc) (n = 14,858 cells). Cell populations were annotated according to canonical marker-gene expression profiles. **(C–D)** Violin plots showing representative marker-gene expression across identified cell clusters in COPD (C) and SSc (D) lung tissues. **(E)** Summary table showing the absolute numbers and relative proportions of cells within each annotated cluster in COPD and SSc lung samples.

**Supplementary Figure 2**

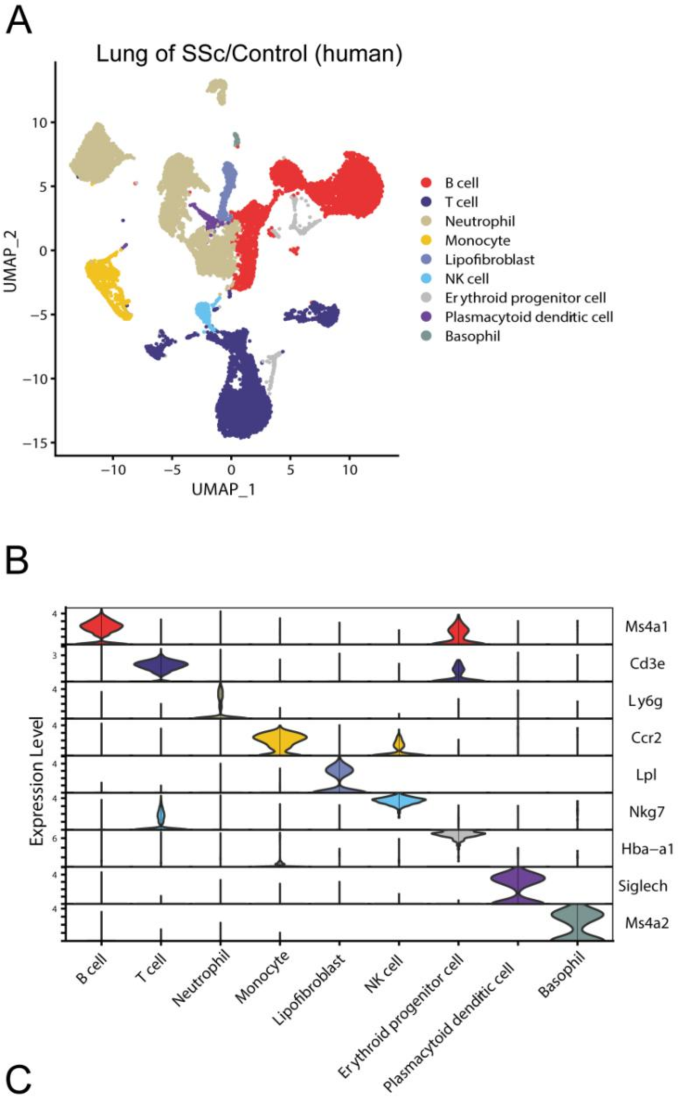


C

| cluster Name | Cell number | | % of total | |
|---|---|---|---|---|
| | Control | COPD | Control | COPD |
| B cell | 4400 | 3228 | 32.43% | 29.32% |
| Neutrophil | 4235 | 3161 | 31.22% | 28.71% |
| Monocyte | 752 | 620 | 5.54% | 5.63% |
| Plasmacytoid dendritic cell | 136 | 113 | 1.00% | 1.03% |
| T cell | 3236 | 3240 | 23.85% | 29.43% |
| Basophil | 68 | 40 | 0.50% | 0.36% |
| Plasmacytoid dendritic cell | 136 | 113 | 1.00% | 1.03% |
| Erythroid progeniter cell | 204 | 145 | 1.50% | 1.32% |
| Lipofibroblast | 400 | 349 | 2.95% | 3.17% |
| total | 13567 | 11009 | | |

**Supplementary Figure 2. Single-cell transcriptomic landscape of peripheral blood from COPD mice. (A)** Uniform Manifold Approximation and Projection (UMAP) visualization of 24,576 peripheral blood cells derived from COPD mice, coloured according to annotated cell subsets. **(B)** Violin plots showing representative marker-gene expression across identified cell clusters. **(C)** Summary table showing the absolute numbers and relative proportions of cells within each annotated cluster.

**Supplementary Figure 3**

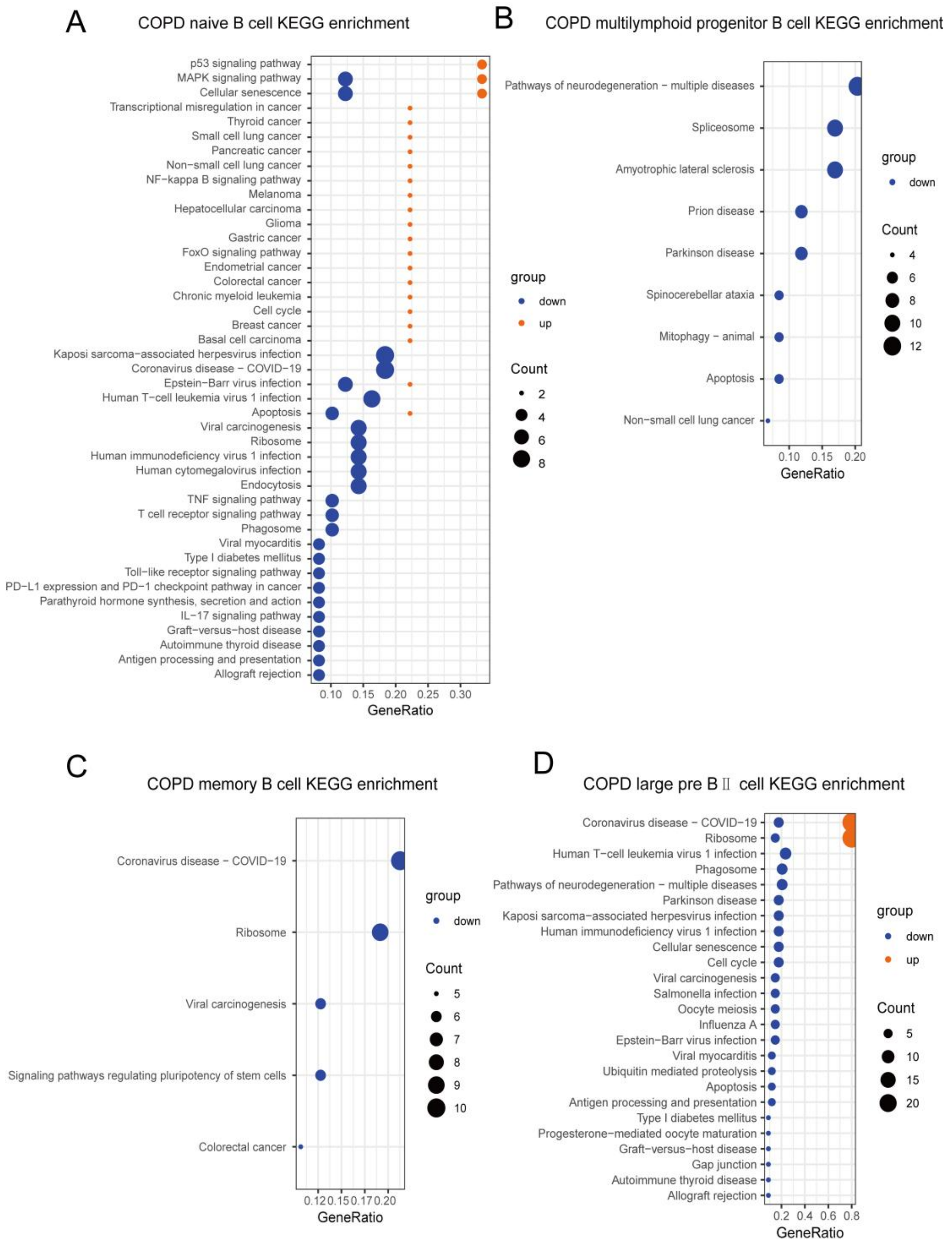


**Supplementary Figure 3. KEGG pathway enrichment analysis of B-cell subsets in the peripheral blood and bone marrow of COPD mice.** Kyoto Encyclopedia of Genes and Genomes (KEGG) pathway enrichment analyses were performed using enrichKEGG for upregulated (red) and downregulated (blue) differentially expressed genes (DEGs) in naïve B cells (A), multi-lymphoid progenitor cells (B) and memory B cells (C) from peripheral blood, as well as large pre-B II cells (D) from bone marrow. Circle size represents the number of genes enriched within each pathway.

**Supplementary Figure 4**

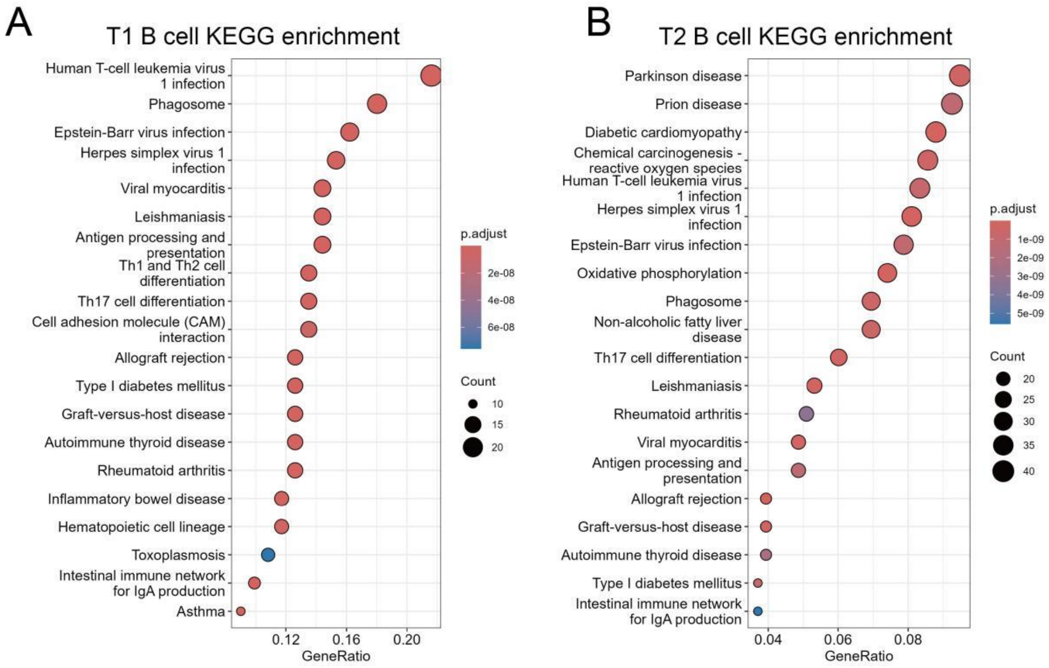


**Supplementary Figure 4. KEGG pathway enrichment analysis of B-cell subsets in the peripheral blood of COPD patients.** Kyoto Encyclopedia of Genes and Genomes (KEGG) pathway enrichment analyses were performed using enrichKEGG for upregulated (red) and downregulated (blue) differentially expressed genes (DEGs) in transitional 1 (T1) B cells (A), and T2 B cells (B).

**Supplementary Figure 5**

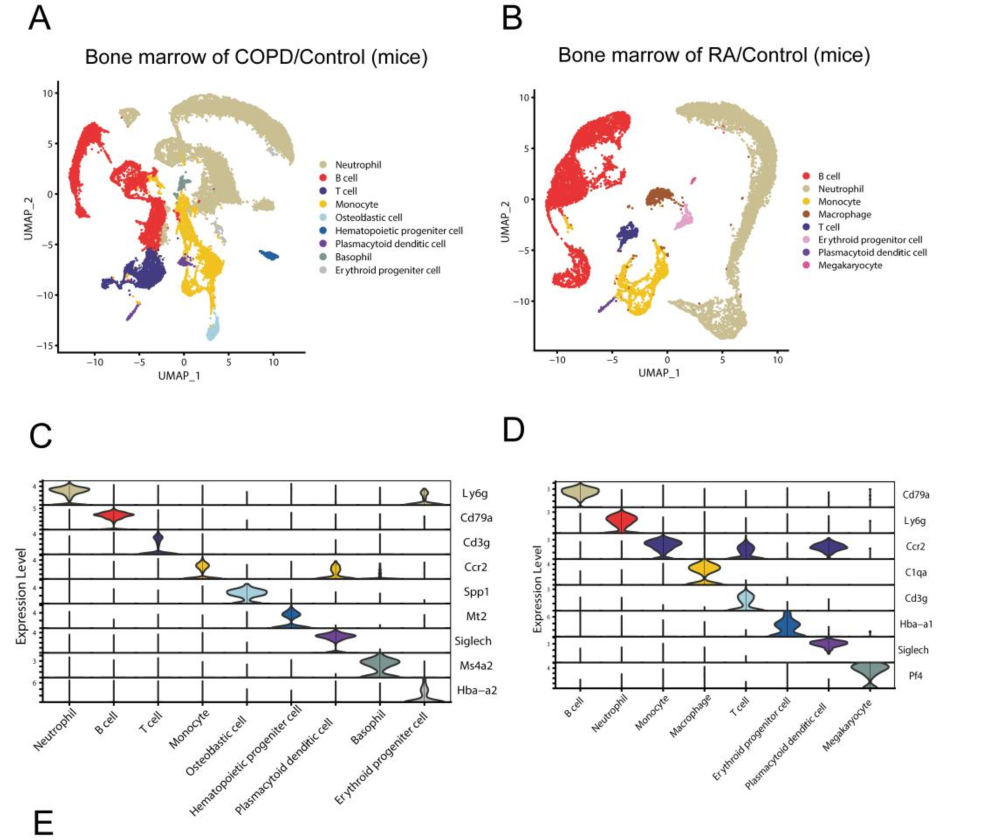


E

| cluster Name | Cell number | | | | % of total | | | |
|---|---|---|---|---|---|---|---|---|
| | Control | COPD | Control | RA | Control | COPD | Control | RA |
| B cell | 3707 | 2816 | 2863 | 1830 | 23.02% | 25.47% | 38.72% | 24.74% |
| Neutrophil | 6563 | 5781 | 3181 | 4038 | 40.76% | 52.29% | 43.02% | 54.60% |
| Macrophage | 0 | 0 | 222 | 388 | 0.00% | 0.00% | 3.00% | 5.25% |
| Monocyte | 2220 | 1347 | 648 | 719 | 13.79% | 12.18% | 8.76% | 9.72% |
| Plasmacytoid dendritic cell | 287 | 229 | 110 | 87 | 1.78% | 2.07% | 1.49% | 1.18% |
| T cell | 2289 | 395 | 224 | 193 | 14.22% | 3.57% | 3.03% | 2.61% |
| Basophil | 139 | 140 | 0 | 0 | 0.86% | 1.27% | 0.00% | 0.00% |
| Osteoblastic cell | 307 | 283 | 0 | 0 | 1.91% | 2.56% | 0.00% | 0.00% |
| Hematopoietic progeniter cell | 476 | 5 | 0 | 0 | 2.96% | 0.05% | 0.00% | 0.00% |
| Erythroid progeniter cell | 112 | 60 | 124 | 126 | 0.70% | 0.54% | 1.68% | 1.70% |
| Megakaryocyte | 0 | 0 | 23 | 15 | 0.00% | 0.00% | 0.31% | 0.20% |
| total | 16100 | 11056 | 7395 | 7396 | | | | |

**Supplementary Figure 5. Single-cell transcriptomic landscape of bone marrow cells from COPD and rheumatoid arthritis (RA) mice. (A–B)** Uniform Manifold Approximation and Projection (UMAP) visualization of bone marrow cells derived from COPD (n = 27,156) and rheumatoid arthritis (RA) (n = 14,791) mice, coloured according to annotated cell subsets. **(C–D)** Violin plots showing representative marker-gene expression across identified cell clusters in COPD (C) and RA (D) mice. **(E)** Summary table showing the absolute numbers and relative proportions of cells within each annotated cluster.

**Supplementary Figure 6**

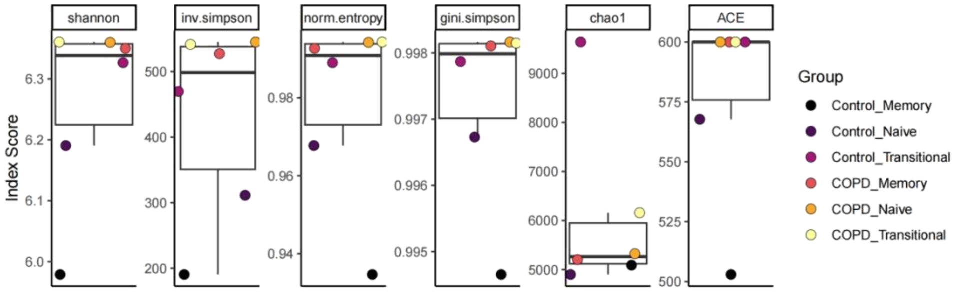


**Supplementary Figure 6. Increased BCR diversity and repertoire richness in COPD B cells.** Compared with healthy controls, B cells from patients with COPD exhibited significantly increased BCR diversity, including higher Shannon entropy, inverse Simpson index and Gini–Simpson index values, together with increased repertoire richness estimates, including Chao1 and ACE scores.

**Supplementary Figure 7**

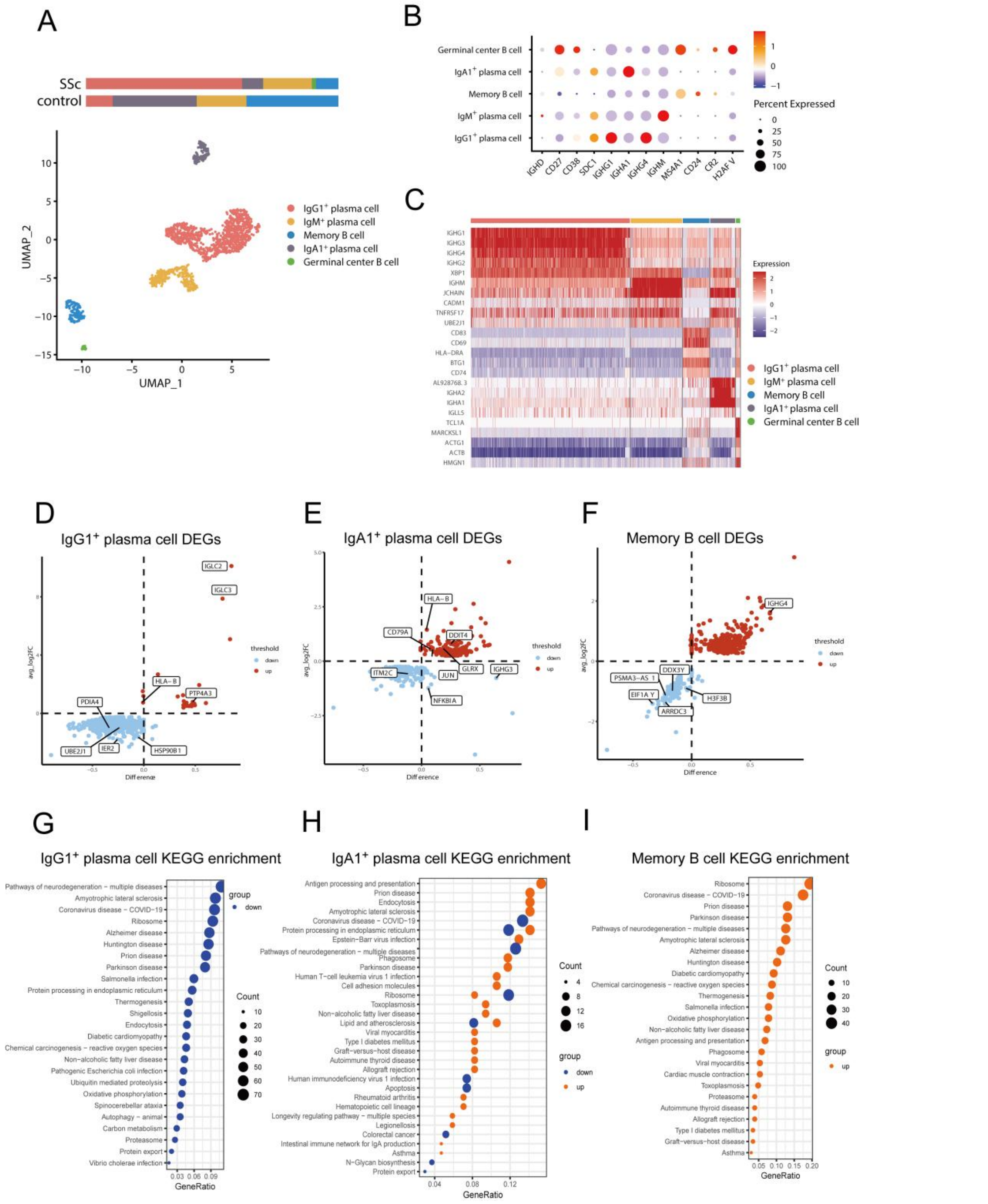


**Supplementary Figure 7. Characteristics of B-cell subclusters in the lungs of patients with systemic sclerosis (SSc). (A)** Uniform Manifold Approximation and Projection (UMAP) visualization of B cells derived from the lungs of patients with systemic sclerosis (SSc) and healthy controls (total n = 1,518 cells), coloured according to annotated B-cell subsets, with relative proportions of each subset indicated. **(B)** Dot plot showing representative marker-gene expression across B-cell subsets. **(C)** Heatmap displaying transcriptional features of individual B-cell subsets. **(D–F)** Volcano plots showing differentially expressed genes (DEGs) in IgG1$^+$ plasma cells (D), IgA1$^+$ plasma cells (E), and memory B cells. (F). Upregulated and downregulated genes are shown in red and blue, respectively. **(G–I)** Kyoto

Encyclopedia of Genes and Genomes (KEGG) pathway enrichment analyses of upregulated and downregulated pathways in IgG1$^{+}$ plasma cells (G), IgA1$^{+}$ plasma cells (H), and memory B cells (I). Upregulated and downregulated pathways are shown in red and blue, respectively. Black circle size represents the number of genes enriched within each pathway.

**Supplementary Figure 8**

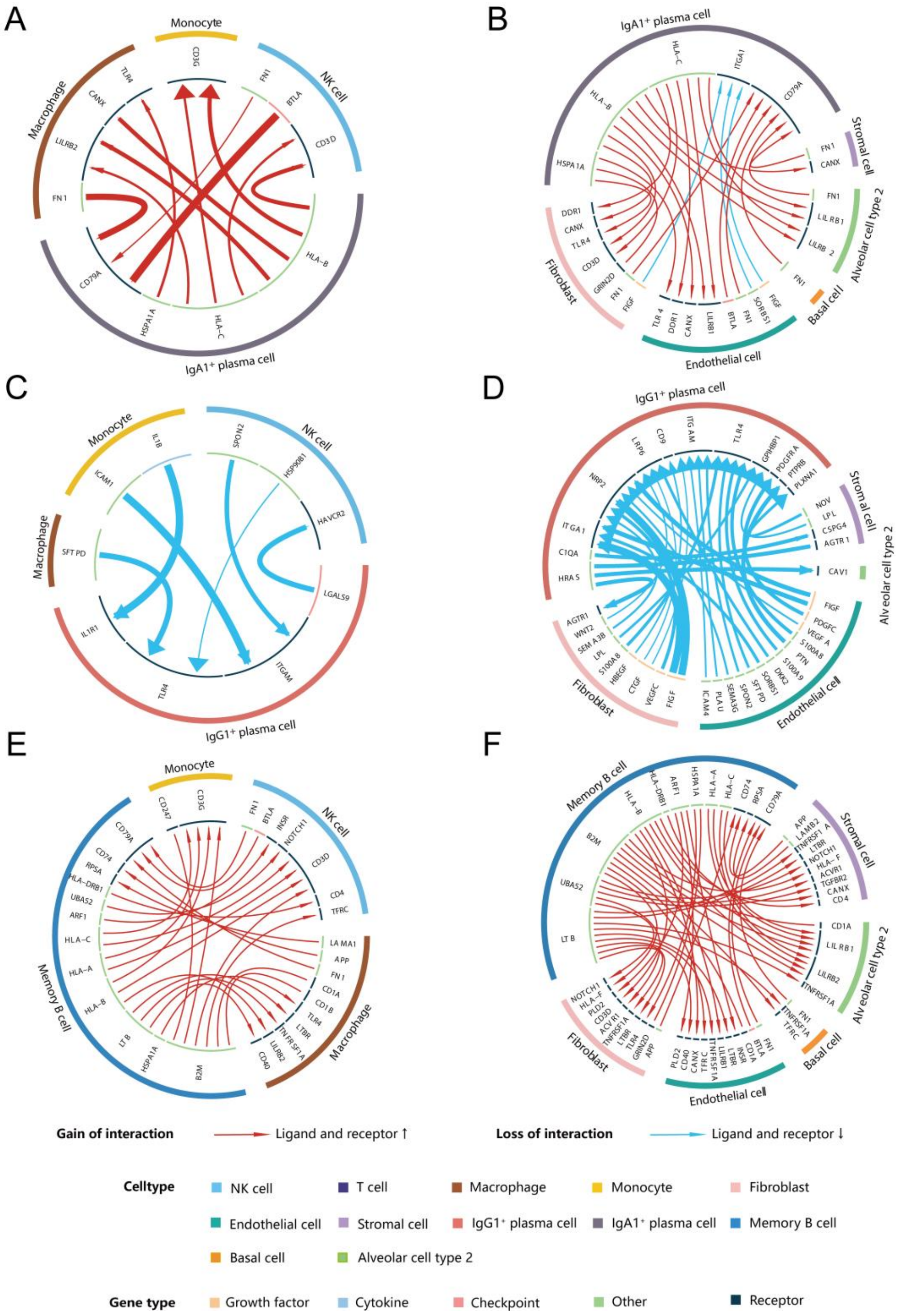


**Supplementary Figure 8. Cell–cell communication analysis of B-cell subsets in the lungs of patients with systemic sclerosis (SSc).** Cell–cell communication analyses were performed to investigate predicted interactions between IgA1$^{+}$ plasma cells (A–B), IgG1$^{+}$ plasma cells (C–D), and memory B cells (E–F) with surrounding immune and non-immune cell populations in the lungs of patients with systemic sclerosis (SSc).

**Supplementary Figure 9**

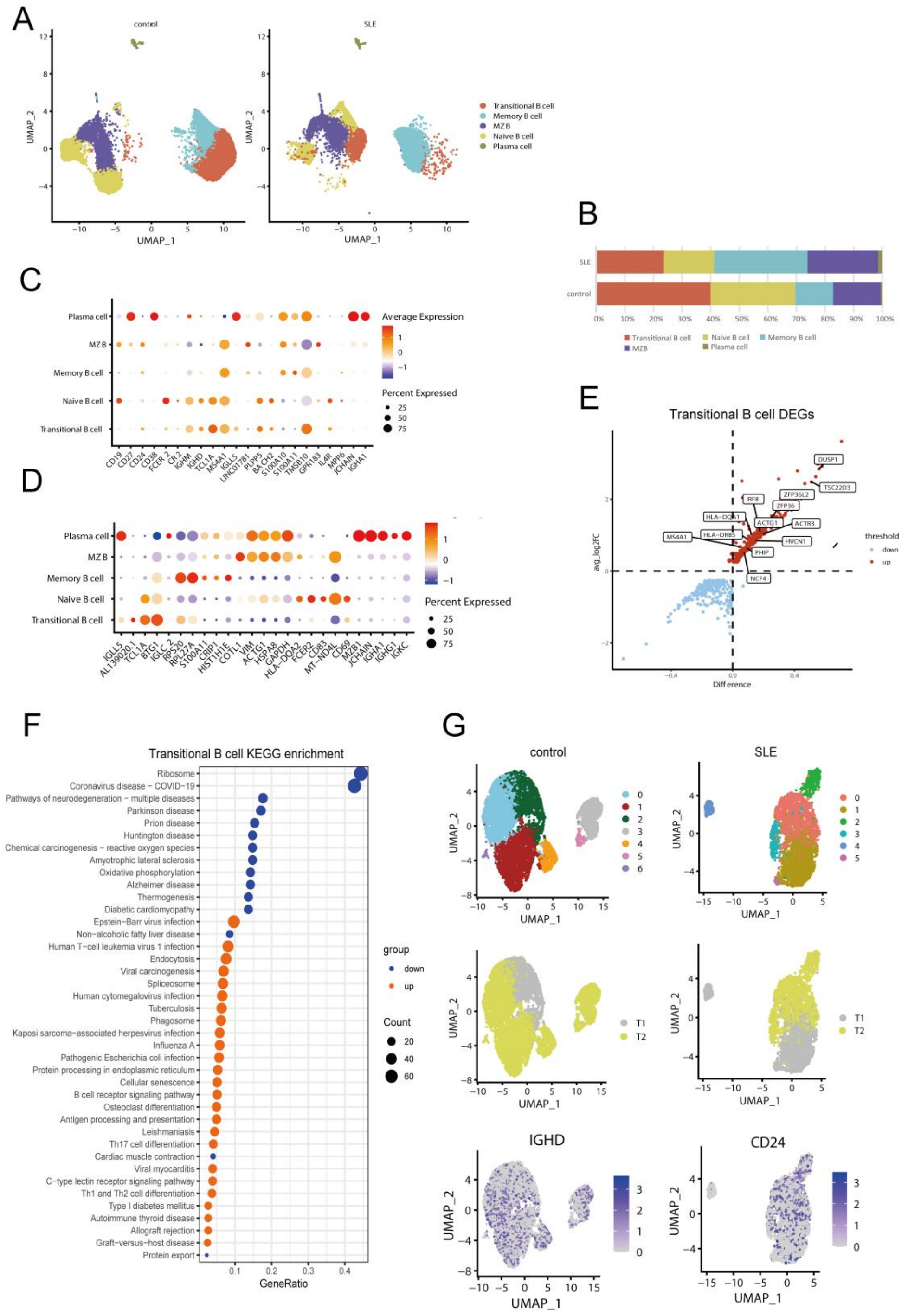


**Supplementary Figure 9. Characteristics of B-cell subclusters in the peripheral blood of patients with systemic lupus erythematosus (SLE). (A–B)** Uniform Manifold Approximation and Projection (UMAP) visualization (A) and relative proportions (B) of B-cell subclusters identified from the peripheral blood of patients with systemic lupus erythematosus (SLE), coloured according to annotated subsets. **(C–D)** Dot plots showing the expression of representative functional marker genes (C) and the top five marker genes (D) across plasma cells, marginal zone B cells (MZBs), activated naïve B cells, naïve B cells, and transitional B cells. **(E)** Volcano plot showing differentially expressed genes (DEGs) in transitional B cells. Upregulated and downregulated genes are shown in red and blue, respectively. Shared DEGs between COPD and SLE are highlighted. **(F)** Kyoto Encyclopedia of Genes and Genomes (KEGG) pathway enrichment analysis of upregulated and downregulated pathways in transitional B cells. Upregulated and downregulated pathways are shown in red and blue, respectively. Black circle size represents the number of genes enriched within each pathway. **(G)** Uniform Manifold Approximation and Projection (UMAP) visualization of transitional B cells from patients with SLE and healthy controls, coloured according to automatically classified clusters and developmental subsets. Expression patterns of classification markers, including IGHD and CD24, are shown for both SLE and control groups.

### Supplementary Figure 10

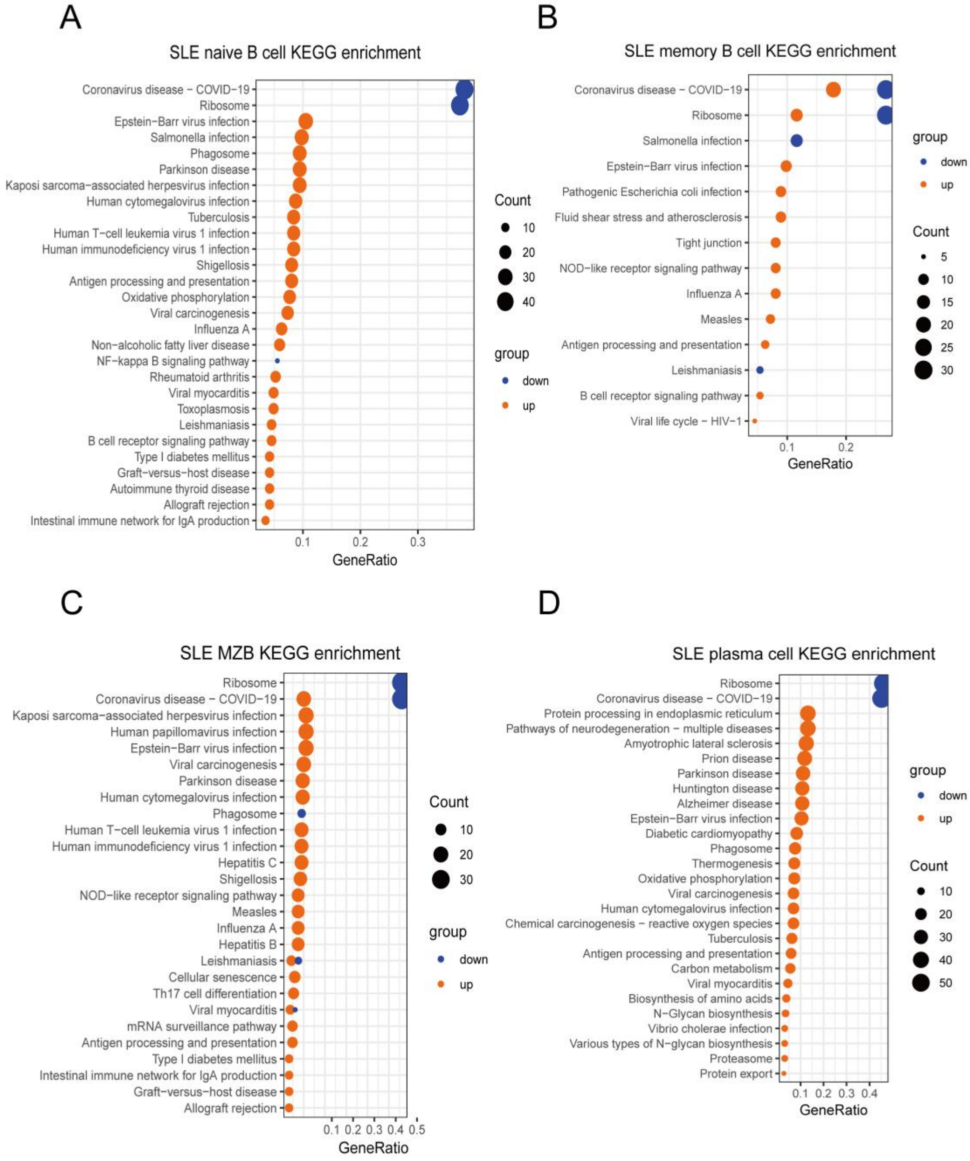


**Supplementary Figure 10. KEGG pathway enrichment analysis of peripheral blood B-cell subsets in patients with systemic lupus erythematosus (SLE). (A–D)** Kyoto Encyclopedia of Genes and Genomes (KEGG) pathway enrichment analyses of upregulated and downregulated pathways among differentially expressed genes (DEGs) in naïve B cells (A), memory B cells (B), marginal zone B cells (C), and plasma cells (D) derived from the peripheral blood of patients with systemic lupus erythematosus (SLE). Upregulated and downregulated pathways are shown in red and blue, respectively. Black circle size represents the number of genes enriched within each pathway.

#### Supplementary Figure 11

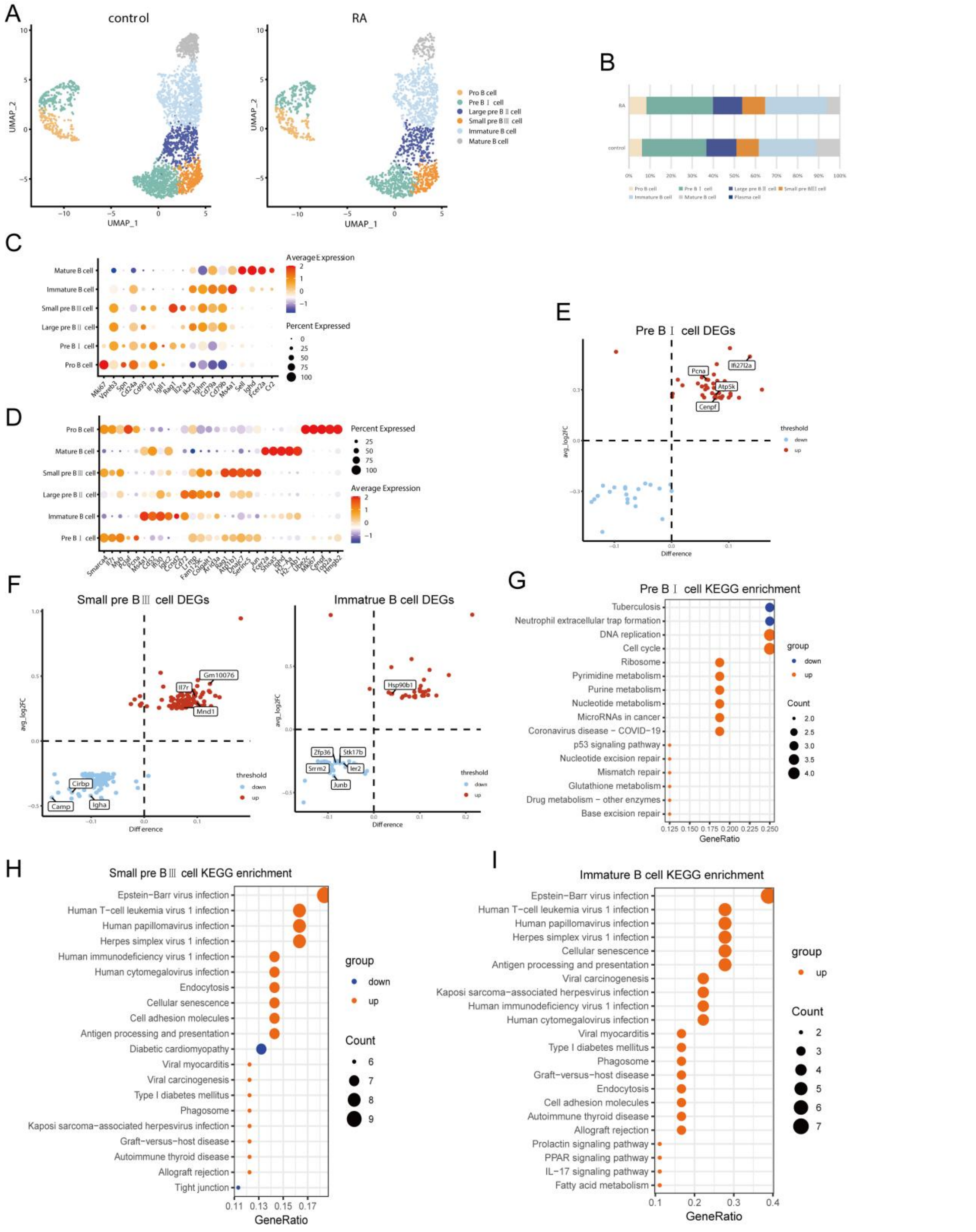


**Supplementary Figure 11. Characteristics of B-cell subclusters in the bone marrow of rheumatoid arthritis (RA) mice. (A–B)** Uniform Manifold Approximation and Projection (UMAP) visualization (A) and relative proportions (B) of B-cell subclusters identified from the bone marrow of rheumatoid arthritis (RA) mice, coloured according to annotated subsets. **(C–D)** Dot plots showing the expression of representative functional marker genes (C) and the top five marker genes (D) across pro-B cells, pre-B I cells, large pre-B II cells, small pre-B III cells, immature B cells, and mature B cells. **(E–G)** Volcano plots showing differentially expressed genes (DEGs) in pre-B I cells (E), small pre-B III cells (F) and immature B cells (G). Upregulated and downregulated genes are shown in red and blue, respectively. Shared DEGs between COPD and RA mice are highlighted. **(H–J)** Kyoto Encyclopedia of Genes and Genomes (KEGG) pathway enrichment analyses of upregulated and downregulated pathways in pre-B I cells (H), small pre-B III cells (I) and immature B cells (J). Upregulated and downregulated pathways are shown in red and blue, respectively. Black circle size represents the number of genes enriched within each pathway.

#### Supplementary Tables

**Supplementary Table S1. Information concerning scRNA-seq data from GEO.**

| Datasets | Organism | Sample source | Platforms | Information concerning models or patients |
|---|---|---|---|---|
| GSE196638 | Homo sapiens | Lung cells from the distal fragments | GPL24676 Illumina NovaSeq 6000 (Homo sapiens) | The normal lung species were obtained from brain-dead donors that were rejected for lung transplantation, while COPD/emphysema lung specimens were taken from the patients undergoing lung transplantation for COPD with radiographic evidence of severe emphysema [1]. |
| GSE205078 | Mus musculus | $CD45^+$ whole blood and $CD45^+$ bone marrow cells | GPL24247 Illumina NovaSeq 6000 (Mus musculus) | Mice were exposed to normal air or cigarette smoke through custom-designed and purpose-built nose-only exposure system. The smoke of up to twelve 3R4F reference cigarettes was administered twice per day, 5 days a week for 12 weeks [2]. |
| GSE132771 | Homo sapiens | Lung cells | GPL24676 Illumina NovaSeq 6000 (Homo sapiens) | Fibrotic lung tissues were obtained at the time of lung transplantation from patients with a diagnosis of scleroderma, while normal lung tissues were obtained from lungs rejected for transplantation by the Northern California Transplant Donor Network [3]. |
| GSE193867 | Homo sapiens | $CD19^+$ peripheral blood mononuclear cells | GPL16791 Illumina HiSeq 2500 (Homo sapiens) | SLE patients samples were obtained with detection of some auto-antibodies, such as ANA, dsDNA, Sm, RNP, C1Q and La [4]. |
| GSE221704 | Mus musculus | Mononuclear cells from bone marrow of femur | GPL24247 Illumina NovaSeq 6000 (Mus musculus) | The collagen-induced arthritis (CIA) model was established by injecting chicken type II collagen subcutaneously into the mouse tail of female DBA/1J mice at the 6th week of age [5]. |

**Supplementary Table S2**
**Primers used for qPCR**

| | Forward primer | Reverse primer |
|---|---|---|
| Igk | TCCATCTTCCCACCATCCAG | GATGTCTTGTGAGTGGCCTC |
| λ5 | TGTGAAGTTCTCCTCCTGCTG | ACCACCAAAGTACCTGGGTAG |
| λ5 promoter | GGGTTAAGACAGGCAGCTGTGAG | CAAACCCCAGGCTGTCTCTAGTT |
| VpreB promoter | TGCCAAGCTGGCCATGTGAACAC | GATGTTCCTCTACCATATGTGAG |
| IL-7R | GGGGCTCTTTTACGAGTGAAATGC | TGTGAGTCTGAGGTAGATGGCCTGC |
| β-actin | AAGACCTCTATGCCAACAC | TAGGAGCCAGAGCAGTAAT |

**Supplementary Table 3. Cellular communication between B cell subsets and other cells in the lung of COPD patients.**

| | Classification | T cells | NK cells | Monocyte-macrophages | Fibroblast | Endothelial | Stromal cells | Basal cells | Alveolar cells |
|---|---|---|---|---|---|---|---|---|---|
| IgA1$^+$ plasma cells | GFs | TGFB1-ACVRL1/CAV1/CD109/TGFBR2/CXCR4↑ | TGFB1-CAV1/CXCR4/TGFBR2/TGFBR3↑<br>FGF2/COL14A1/HBEGF/HAS2-CD44↑ | TGFB1-TGFBR3/ITGB8/CD109/SDC2/ITGAV↑<br>FGF2/HBEGF/-CD44↑ | TGFB1-CXCR4/ITGB8↑<br>FGF1/FGF2/HGF/HBEGF-CD44↑ | TGFB1-CXCR4/TGFBR1/ITGAV/ACVRL1↑<br>FGF2-SDC1↓ | TGFB1-CXCR4/ENG/SDC2/TGFBR2↑ | TGFB1-ITGB6↑<br>HBEGF-CD44↑ | TGFB1-CXCR4/ITGB8/CD44/ACVRL1/ENG/ITGAV↑<br>HBEGF-CD44↑ |
| | CKs | IL1B-IL1R1↑<br>IL6-IL6R↑ | IL1B-IL1R1↑<br>IL6-IL6R↑ | IL1B/IL1A/IL1RN-IL1R1↑ | IL1RN/IL1B-IL1R1↑<br>IL6-IL6R↑ | IL1RN-IL1R1↑<br>IL6-IL6R↑ | IL1B-IL1R1↑<br>IL6-IL6R↑ | IL1B-IL1R1↑<br>IL6-IL6R↑ | IL1B/IL1RN-IL1R1/IL1RAP↑<br>IL6-IL6R↑ |
| | Chemokines | CCL20-CCR6↑<br>CCL7-CCR2↓<br>CCL13-CCR2/5↓ | CCL7/13-CCR1/2↓ | XCL2-XCR1↑<br>CCL20-CCR6/CXCR3↑<br>CCL13-CCR5↓<br>CCL13-CCR1/2↓ | CCL2-ACKR4↑<br>CCL20-CXCR3↑<br>CCL13-CCR2↓ | CCL2/3/5-CCR4↑<br>CCL20-CCR6↑ | CCL2-CCR10↑<br>CCL5-SDC4↑ | CCL5-SDC4↑ | CCL5-GPR75↑ |
| | Checkpoints | ICOSLG-ICOS↑ | ICOSLG-ICOS↑<br>CD40LG-TRAF3↑ | ICOSLG-ICOS↑<br>CD40LG-TRAF3↑ | ICOSLG-ICOS↑<br>CD40LG-TRAF3↑ | ICOSLG-ICOS↑ | ICOSLG-ICOS↑ | | ICOSLG-ICOS↑ |
| | Receptors | CD70-CD27↓<br>SELPLG-SELL↓<br>SPP1-S1PR1/ITGB1↓ | | PTGS2-PTGDR2↑<br>HLA-A/HLA-B-KIR2DL3↑<br>WNT4-FZD2↓ | | HLA-A-KIR3DL2↑ | | | HLA-A-KIR3DL2↑<br>WNT5A-FZD2 ↓ |
| | Others | PTGS2-CAV1↑<br>SERPING1-SELE↑<br>GPC3-CD81↑ | PTGS2-CAV1↑<br>SERPING1-SELE↑<br>FN1-C5AR1/CD79A↑ | SERPING1-SELE↑<br>FN1-COL13A1↑<br>BTLA-CD79A↑<br>SEMA6D-TYROBP↑<br>HDC-HRH2↑<br>COL18A1-GPC1↑<br>CALM1/2-MYLK↑<br>HGF-SDC1↓<br>SPP1-ITGA9↓ | HDC-HRH1↑<br>THBS1-ITGB3/SDC4/ITGA6/ITGA4/SDC1↑ | FN1-MAG↑<br>C3-CD19↑<br>HDC-HRH1↑<br>THBS1-TNFRSF11B/ITGB3↑ | FN1-CD79A↑<br>HGF/HBEGF/-CD44↑<br>HDC-HRH2↑<br>PTGS2-ALOX5↑<br>THBS1-ITGA4/LRP1↑<br>RARRES2-CMKLR1↑ | HDC-HRH1↑<br>THBS1-ITGA6/ITGB1/SDC4/CD47↑<br>FBLN1-ITGB1↑ | HDC-HRH1/2↑<br>SELPLG-SELE↑<br>THBS1-ITGA4↑ |
| IgG1$^+$ plasma cells | GFs | | FGF2/HBEGF/COL14A1/HAS2-CD44↑ | HBEGF/FGF2/COL14A1-CD44↑ | FGF1/2/HGF/HBEGF-CD44↑ | | HGF/HBEGF-CD44↑ | HBEGF-CD44↑ | HBEGF-CD44↑ |
| | CKs | IL1B-IL1R1↑<br>IL6-IL6R↑ | IL1B-IL1R1↑<br>IL6-IL6R↑ | IL1B-IL1R1/IL1R2/IL1RAP↑<br>IL6-IL6R↑ | IL1B-IL1R1/IL1RAP↑<br>IL6-IL6R↑ | IL1B-ADRB2/IL1RAP/IL1R1↑<br>IL6-IL6R↑ | IL1B-IL1R1↑<br>IL6-IL6R↑ | IL1B-IL1R1↑<br>IL6-IL6R↑ | IL1B-IL1R/IL1RAP/ADRB2↑<br>ICAM1-IL2RA↑<br>IL6-IL6R↑ |
| | Chemokines | | | CCL2/3/4-CCR5↑<br>CXCL2-XCR1↑ | CCL2-ACKR4↑ | CCL2/3-CCR4↑ | CCL2-CCR10↑ | | |
| | Checkpoints | | CD40LG-TRAF3↑ | CD40LG-TRAF3↑ | CD40LG-TRAF3↑ | | | | |

| | | | | | | | | | |
|---|---|---|---|---|---|---|---|---|---|
| | Receptors | | | HLA-B-KIR2DL3↑ | | | | | |
| | Others | SERPING1-SELE↑<br>FN1-C5AR1↑<br>COL14A1-CD44↑<br>COL6A2-ITGA1↑ | SERPING1-SELP↑<br>CALM1/2-MYLK↑<br>VWF-SELP↑ | SERPING1-SELE↑<br>CALM1-MYLK/IL2RA↑<br>FN1/BTLA-COL13A1/SDC2/CD79A↑<br>SCGB1A1-LMBR1L↑ | CALM1/2/3-MYLK↑<br>THBS1-ITGB3/SDC4↑<br>DCN-ERBB4↑ | THBS1-TNFRSF11B/ITGB3↑<br>CALM2-MYLK↑<br>FN1-MAG↑<br>ICAM1-IL2RA↑<br>VWF-TNFRSF11B↑ | SERPING1-SELE↑<br>CALM1/2/3-MYLK↑<br>FN1-CD79A↑<br>THBS1-ITGA4↑ | CALM2-MYLK↑<br>COL6A2/THBS1-ITGA6/ITGB1↑<br>ITGA6/ITGB6↑<br>-ITGB1↑ | CALM1/2-MYLK↑<br>COL6A2-ITGA2↑<br>FN1-ITGA2↑<br>APOE-LRP8↑ |
| **Memory B cells** | GFs | TGFB1/CYR61/ICAM1-CAV1/CXCR4↑ | TGFB1/PTGS2/CYR61-CAV1↑<br>TGFB1/CXCL12-CXCR4↑<br>FGF2/HBEGF/COL14A1/HAS2-CD44↑ | FGF2/HBEGF/-CD44↑ | TGFB1/ICAM1-CAV1/CXCR4↑<br>FGF1/FGF2/HGF/HBEGF-CD44↑ | TGFB1-CAV1/CXCR4↑ | TGFB1/ICAM1-CAV1↑<br>TGFB1/CXCL12-CXCR4↑<br>HGF/HBEGF/-CD44↑ | HBEGF-CD44↑ | HBEGF-CD44↑<br>TGFB1-CAV1↑<br>HBEGF-CD44↑<br>TGFB1-CXCR4↑ |
| | CKs | IL1B-IL1R1↑<br>IL10-IL10RA↑<br>TNFRSF14/CD247-BTLA↓ | IL1B-IL1R1↑<br>TNFRSF14-BTLA↓ | IL1B-IL1R1/IL1R2↑ | | IL1B-ADRB2↑ | IL1B-IL1R1↑<br>ICAM1-IL2RA↑ | IL1B-IL1R1↑ | IL1B-IL1R1/IL1RAP↑ |
| | Chemokines | CCL21-CCR7↑<br>CXCL12-CXCR4↑<br>CCL18-C14orf1↓ | CCL19/CCL21-CCR7↑<br>CCL18-C14orf1↓ | CCL19/CCL21-CCR7/CXCR3↑<br>CCL3L3/CCL2/CCL4-CCR5↑ | CCL2/CCL21-ACKR4/CCR7/-CXCR3↑<br>CCL5-CXCR3↑ | CCL19/CCL21-ACKR2/CCR7↑<br>CCL2/5-CCR4↑ | CCL21-CCR7↑<br>CCL2-CCR10↑<br>CCL5-SDC4↑ | CCL21-CCR7↑<br>CCL5-SDC4↑ | CCL19/21-CCR7↑<br>CCL5-GPR75↑<br>CCL18-C14orf1↓ |
| | Receptors | CD70-CD27↓ | | HLA/B2M-A-KIR2DL3↑ | CD70-CD27↑<br>NCR3LG1-NCR3↑ | HLA-A-KIR3DL2↑<br>NCR3LG1-NCR3↑ | | | CD70-CD27↑<br>NCR3LG1-NCR3↑<br>BTLA-TNFRSF14↓<br>TNFRSF14-BTLA ↓ |
| | Others | CCL5-SDC4↑<br>ANXA1-FPR2↑<br>COL1A2-ITGA1↑<br>ICAM1-IL2RA↑<br>BMP8B-ACVR1↓ | FN1-CD79A↑<br>COL1A2-CD93↑ | BTLA-CD79A↑<br>FN1-SDC2/ITGA8/COL13A1↑<br>ICAM1-IL2RA↑<br>COL1A2-ITGA2↑<br>BMP8B-ACVR1/2A/2B/BMPR2↓ | TGM2/COL1A2-ITGB3/SDC4/ITGA9↑<br>BMP8B-ACVR2B↓ | FN1-MAG↑<br>COL1A2-CD44↑<br>BMP8B-ACVR1/ACVR2A/BMPR1A↓ | TGM2-ITGA4↑<br>CALM1-SCTR↑ | ICAM1-CAV1↑<br>TGM2-ITGB1↑<br>TGM2-SDC4↑<br>COL1A2-ITGB1↑<br>ICAM1-IL2RA↑<br>FN1-ITGB6↑ | BMP8B-ACVR2A↓<br>BMP8B-BMPR1A ↓<br>HLA-A-KIR3DL2↑<br>COL1A2-ITGA2↑<br>ICAM1-IL2RA↑<br>COL1A2-CD93↑ |

**Supplementary Table 4. The common DEGs between COPD and other autoimmune diseases in bone marrow, peripheral blood and lung.**

| Sample sources | subclusters | common DEGs between COPD and autoimmune diseases | |
|---|---|---|---|
| | | Up | Down |
| lung | IgG1$^{+}$ plasma cell | IGLC2, IGLC3, PTP4A3, HLA-B, RP11-138A9.2 | IER2,SUB1,PPIB,HSP90B1,UBE2J1,TMED9,TPD52,TNFRSF17,COX5B,RPS26,RSRP1,PPIA,SRP14,EGR1,PDIA4,TAF7,MYDGF,SEC61G,COPE,SPCS1,UBXN4,QPCT,PRDX1,UQCR10,EDF1,SON,SPCS2,MTCO1,C11orf31,GOLGB1,RPS4Y1,BMI1,DUSP6,EAF2,SEC62,PSAP,SMARCB1,LRRC16A,ATP5H,HSH2D,C14orf2,TMEM208,USO1,ZNHIT1,TMED2,NDUFB3,TMEM134,C19orf60,C1QB,CD2BP2,MTND4,MPHOSPH8,ANAPC5,ACIN1,TRAPPC2L,MRPL57,AKAP17A,NDUFA4,KLF4,IL2RG,KATNBL1,MZB1,SMC1A,CCDC167,IK,PFDN6,VPS36,DOCK8,ANXA6,B9D1,GOLGA8B,CD27,SDF4,SESN1,BTN3A2,ILF3-AS1,EDEM3,ATP5E |
| | IgA1$^{+}$ plasma cell | HLA-B, RPL17, HLA-DPA1, DDIT4, IGKV3-20, GLRX, RPS10, CD79A, RPL41, H2AFZ, HLA-A, CHST2, BASP1, PRDM1 | NFKBIA, RPS26, IGHG3, JUN, ITM2C, BRD2, ARGLU1, AMPD1, BCL2, RPN2, PAIP2B, HNRNPA2B1 |
| | Memory B cell | IGHG4, RPL17, RPL36A, RPS10, RP11-138A9.1, PTGES3, RPS21, HLA-DPB1, HLA-DQA1, RPL41 | RPS4Y1, EIF1AY,ARRDC3, N4BP2L2, LINC00969, PSMA3-AS1, ASH1L, AES, DDX3Y, H3F3B, PNISR, SFTPB, ATF7IP2, LRRK2, LRRC16A, SLPI |
| Peripheral blood | Transitional B cell | Zfp36l2,Tsc22d3,Ms4a1,Dusp1,Zfp36,Actr3,Ncf4,Actg1,Irf8,Napsa,Phip,Rgs19,Hvcn1,H2-Aa,H2-Eb1 | Rpl37,Rps29,Rps27,Rpl38,Rpl35,Rps28,Rpl27,Rpl29,Uba52 |
| | Resting naive B cell | Dusp1,Gadd45b | Gdi2,Pnrc1,Arf6,Id3,Rpl27,mt-Nd4l,Rpl29 |
| | Memory B cell | Tsc22d3,March1,Irf8,Ms4a1,Bdp1,Pou2f2,Chordc1 | Rps28,H3f3b,Jchain |
| | Plasma cell | Arf1,Gapdh,Ergic2,Wipi1,Bloc1s2,Trabd,Spn,Uqcr10 | None |
| Bone marrow | Pro B cells | None | Tmsb10, Cirbp |
| | Pre B I cells | Ifi27l2a, Pcna, Atp5k, Cenpf | None |
| | Large pre B II cells | Mnd1, Gm10076 | Cirbp, Pafah1b3, Igha |
| | Small pre B III cells | Il7r、Mnd1、Gm10076 | Cirbp、Camp、Igha |
| | Immature B cells | Hsp90b1 | Zfp36,Ier2,Stk17b,Srrm2,Junb |
| | Mature B cells | Ptpn22、Ddit4、Zfp36l2、Pou2af1、Ccnd3、Kmt2e、Scd1、Tnfrsf13c、Mybbp1a、Dnaja1 | Nup210,B4galnt1,Clec2i,Tomm6,Zfp36l1 |

**Supplementary Table 5. Cell communication between B subclusters with other cells in SSc patients.**

| B subclusters | Classification | NK cell | Mono-macropage | Fibroblast | Endothelials | Stromal cells |
|---|---|---|---|---|---|---|
| IgA1$^{+}$ plasma cells | Receptors | HLA-B/C-CD3D↑ | HSPA1A-TLR4↑ HLA-B/C-CD3G/LILRB2↑ | HSPA1A-TLR4↑ HLA-B/C-CD3D↑ | HSPA1A-TLR4↑ HLA-B/C-LILRB1↑ | |
| | Others | BTLA/FN1-CD79A↑ | FN1-CD79A↑ HLA-B-CANX↑ | FN1-CD79A↑ HLA-B-CANX↑ HLA-C-DDR1↑ FIGF-ITGA1↓ HSPA1A-GRIN2D↑ | BTLA/FN1-CD79A↑ HLA-C-DDR1↑ HLA-B-CANX↑ FIGF/SORBS1-ITGA1↓ | FN1-CD79A↑ HLA-B-CANX↑ |
| IgG1$^{+}$ plasma cells | GFs | | | VEGFC-NRP2↓ | VEGFA-NRP2↓ PDGFC-PDGFRA↓ | |
| | CKs | | IL1B-IL1R1↓ | | | |
| | Receptors | HSP90B1-TLR4↓ LGALS9- | SFTPD-TLR4↓ ICAM1-ITGAM↓ | WNT2-LRP6↓ S100A8-TLR4↓ | DKK2-LRP6↓ S100A8/S100A9/SFTPD-TLR4↓ | |

| | | | | | | |
|---|---|---|---|---|---|---|
| | | HAVCR2↓ | | | ICAM4-ITGAM↓ | |
| | Others | SPON2-ITGAM↓ | | CTGF-LRP6/ITGAM↓<br>FIGF-ITGA1↓<br>HBEGF-CD9↓<br>LPL-GPIHBP1↓<br>HRAS-AGTR1↓<br>FIGF/SEMA3B-NRP2↓ | FIGF/SEMA3G-NRP2↓<br>FIGF-ITGA1↓<br>PTN-PTPRB↓<br>SORBS1-ITGA1↓<br>SPON2/PLAU-ITGAM↓ | HRAS-AGTR1↓<br>C1QA-CSPG4↓<br>LPL-GPIHBP1↓<br>NOV-PLXNA1↓ |
| Memory B cell | CKs | | LTB-LTBR/TNFRSF1A/CD40↑ | LTB-LTBR/TNFRSF1A↑ | LTB-LTBR/TNFRSF1A/CD40↑ | LTB-LTBR/TNFRSF1A↑ |
| | Receptors | HLA-A/B/C/B2M-CD3D↑<br>HLA-DRB1-CD4↑ | HSPA1A-TLR4↑<br>HLA-A/B/C-CD3G/LILRB2↑<br>B2M-CD3G/CD1A/B/247↑ | HSPA1A-TLR4↑<br>HLA-B-CD3D↑<br>B2M-HLA-F↑ | B2M-CD1A↑<br>HLA-B-LILRB1↑ | B2M-HLA-F↑<br>HLA-DRB1-CD4↑ |
| | Others | ARF1-INSR↑<br>UBA52-NOTCH1↑<br>FN1/BTLA-CD79A↑<br>B2M-TFRC↑ | FN1-CD79A↑<br>APP-CD74↑<br>LAMA1-RPSA↑ | HSPA1A-GRIN2D↑<br>ARF1-PLD2↑<br>UBA52-NOTCH1↑<br>APP-CD74↑<br>UBA52-ACVR1↑ | ARF1-INSR↑<br>FN1/BTLA-CD79A↑<br>ARF1-PLD2↑<br>HLA-B-CANX↑<br>B2M-TFRC↑ | UBA52-ACVR1/NOTCH1/TGFBR2↑<br>APP-CD74↑<br>HLA-B-CANX↑<br>LAMB2-RPSA↑ |
| IgM$^+$ plasma cell | Receptors | HLA-B/C-CD3D↑<br>C1QA-CD93↓ | HSPA1A-TLR4↑<br>HLA-B/C-CD3G/LILRB2↑ | EFNB2-EPHB6↓<br>WNT2-FZD7↓<br>C1QA-CD93↓ | EFNB1-EPHA4↓<br>HLA-B/C-LILRB1↑ | C1QA-CD93↓ |
| | Others | FN1/BTLA-CD79A↑<br>VIM-CD44↑<br>OSM-LIFR↓<br>SFTPA2-CD93↓ | HLA-B-CANX↑<br>VIM-CD44↑<br>FN1-CD79A↑ | HSPA1A-GRIN2D↑<br>SEMA3B-NRP2↓<br>ADM-RAMP2/CALCRL↓<br>SLIT2-ROBO4↓<br>SFTPA2-CD93↓<br>LPL-GPIHBP1↓<br>HBEGF-PRLR↓ | OSM-LIFR↓<br>FN1/BTLA-CD79A↑<br>HLA-C-DDR1↑<br>VIM-CD44↑<br>SFTPA2-CD93↓<br>EFNA5-EPHA4↓<br>PTN-PTPRB↓ | VIM-CD44↑<br>FN1-CD79A↑<br>HLA-B-CANX↑<br>C1QA-CSPG4↓<br>SFTPA2-CD93↓<br>LPL-GPIHBP1↓ |
